# A single large restored patch has lower tree diversity than several smaller ones

**DOI:** 10.1101/2025.11.15.688657

**Authors:** Carmen Galán-Acedo, Federico Riva, Lenore Fahrig, Dirk Hölscher, Amanda E. Martin

**Affiliations:** Department of Biology, Carleton University, Ottawa, Ontario, Canada; Institute for Environmental Studies, Vrije Universiteit Amsterdam, NU building, De Boelelaan 1111, 1081 HV Amsterdam, The Netherlands; Tropical Silviculture and Forest Ecology, University of Goettingen, Büsgenweg 1, 37077 Göttingen, Germany; Centre for Biodiversity and Sustainable Land Use (CBL), University of Goettingen, Büsgenweg 1, 37077 Göttingen, Germany; National Wildlife Research Centre, Environment and Climate Change Canada, Ottawa, Ontario, Canada

**Keywords:** fragmentation, habitat configuration, Kunming-Montreal Global Biodiversity Framework, restoration, oil palm, SLOSS, small patches, species diversity, UN Decade on Ecosystem Restoration

## Abstract

1. Restoration initiatives often target restoring the largest possible amount of habitat to provide the greatest benefits for biodiversity. However, the optimal configuration (e.g., the size and number of restored patches) of habitat, given a fixed total area, remains an unresolved question.

2. Here, we ask whether restoring a single large habitat patch or a mixture of smaller patches of the same total area supports higher plant diversity. To address this question, we measured taxonomic, phylogenetic, and functional diversity of all naturally recruiting woody species in 52 restored vegetation patches in Jambi Province, Sumatra, Indonesia. Thirteen restored patches of each of four sizes (25, 100, 400, and 1,600 m²) were established within conventional oil palm plantations six years before vegetation sampling. From these 52 patches, we generated 750 random comparisons between a single large patch vs. several small patches, ensuring equal total area (100, 400, or 1,600 m²). We evaluated taxonomic, phylogenetic, and functional diversity separately for all species, for native species, and for native forest species, using three diversity measures: species richness, the exponential of Shannon entropy, and the inverse of Simpson concentration.

3. Our findings indicate that restoring several smaller patches results in greater taxonomic, phylogenetic, and functional diversity of recruiting woody species than restoring a single large patch of the same total area. This result holds across the three total habitat areas (100, 400, and 1,600 m²), all species groupings, and all diversity metrics. As expected, species diversity also increased with total area in all cases.

4. Our findings challenge restoration strategies that focus exclusively on enlarging patches. Instead, biodiversity will be enhanced by increasing the total restored area across many patches of different sizes, including very small ones (e.g., 25 m²).

## Introduction

Loss of natural habitats through land use change is the primary threat to biodiversity (Caro et al., 2022; Newbold et al., 2016). In an effort to prevent, halt, and reverse ecosystem degradation, the United Nations (UN) Assembly has formally adopted a resolution designating 2021–2030 as the UN Decade on Ecosystem Restoration (UN-DER; United Nations Environment Programme 2022). National, regional, and global initiatives, such as the Kunming-Montreal Global Biodiversity Framework, emphasize the need for evidence-based restoration policies that effectively target ecosystem recovery (Stephens, 2023). Evidence shows that restoring the largest possible amount of natural habitat promotes biodiversity (Chazdon et al., 2025; Dendy et al., 2015). Yet, the optimal configuration (i.e., the spatial arrangement of habitat such as the sizes and number of patches) of restored habitat for biodiversity, for a given amount of habitat, remains poorly understood (Riva et al., 2025; Watts & Hughes, 2024). This knowledge is particularly needed for restoring key ecosystems such as tropical forests, which have been largely affected by anthropogenic land use such as oil palm plantations (Curtis et al., 2018; Descals et al., 2021), with detrimental effects on tropical biodiversity (Gibson et al., 2011; Struebig et al., 2025).

Recommendations on the configuration of restored habitat have generally focused on reducing fragmentation, by prioritizing actions that connect remnant habitat patches (Antongiovanni et al., 2022; Clark et al., 2023; Gann et al., 2019; Neyret et al., 2025) or increase the area of remnant patches (Banks-Leite et al., 2020; Fink et al., 2009; Watts & Hughes, 2024). However, there is little empirical evidence that these suggested spatial arrangements result in higher biodiversity than more but smaller restored patches, for a given total area of restored habitat. For instance, Watts & Hughes (2024) suggest that ‘*creating more fragmented patches, isolated from source populations, may limit benefits to biodiversity*’, but did not cite any empirical studies showing negative effects of fragmentation on biodiversity in a restoration context (Riva et al., 2025). In contrast to these recommendations, empirical and modelling studies have found that increasing fragmentation (i.e., the number of patches) in a restoration context may benefit biodiversity equally or even more than restoring the same amount of habitat to increase patch sizes or connect patches (Schultz & Crone 2005; Nicol & Possingham 2010; Anonymous, in press).

To make recommendations for the configuration of restored habitat, we need to answer a key question: when does restoring a single large habitat patch or multiple small patches, *for a fixed amount of restored area*, yield better outcomes (Fig. 1)? This question has been extensively studied in the context of biodiversity conservation and protected areas (i.e., Single Large vs. Several Small [SLOSS] debate; Diamond 1975; Ovaskainen 2002; Lindenmayer et al. 2015; Fahrig et al. 2022), but is not well studied in a restoration context. While it was initially suggested that, for the same total habitat area, a single large reserve promotes greater biodiversity than several small reserves (Diamond, 1975), evidence has shown that several small habitat patches usually host as many or more species as a single contiguous habitat patch (Lindenmayer et al., 2015; Quinn & Harrison, 1988; Riva & Fahrig, 2022), though this is still under debate. This could be, in part, because many small patches are spread over a larger total spatial extent than a single large patch, which can lead to higher microhabitat diversity (i.e., greater environmental heterogeneity) and thus greater species richness (Fahrig, 2020). In addition, if the effects of patch area on species occupancy are relatively weak, then a higher number of smaller patches increases the likelihood that a species is present in at least one patch in the landscape.

**Figure 1.**
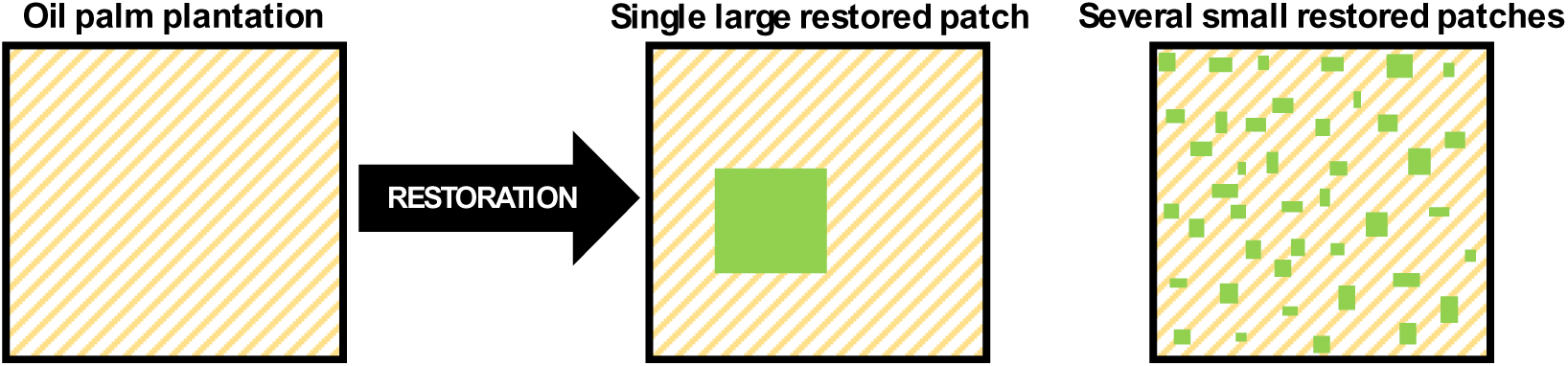
In this hypothetical example, the first square represents a landscape composed of an oil palm plantation. Restoring a particular area of forest could be done in a single large restored forest patch (green rectangle, second landscape) or in many smaller patches of different sizes (green smaller rectangles, third landscape).

Despite the increasing recognition of the cumulative conservation value of small patches over the last decade (Arroyo-Rodríguez et al., 2022; Wintle et al., 2019), there remains a persistent concern that many species might have minimum patch size requirements (Brown & Crone, 2016; Ramirez-Reyes et al., 2016; Riva & Fahrig, 2023b). If so, then these species would not be able to maintain viable populations and finally would be absent from landscapes containing many small patches, even if the total area of habitat in the landscape is large. Such landscapes might therefore only support a deterministic subset of species found in a single large patch. Understanding whether these expectations hold within restored habitat is important for guiding future restoration efforts.

Another common concern about small patches is that the higher biodiversity of several small rather than a single large patch is driven by generalist and invasive species (Fletcher et al. 2018; Chase et al. 2020). Specialist species are thought to be particularly vulnerable to small patches due to their strong dependence on patch-specific resources (Lövei et al., 2006; Miller et al., 2015). In contrast, habitat generalists are suggested to have greater capacity to inhabit and/or disperse through the matrix (i.e., non-habitat areas surrounding habitat patches) (Miller et al., 2015). This leads to the expectation that species richness of habitat specialists and native species is lower in several small patches than a single large one because the composition of larger patches may provide critical habitat for specialists and threatened species. While most studies have found more specialist species across several small than a single large patch (Rösch et al. 2015; Fahrig 2020; Riva & Fahrig 2023a), the opposite has also been found (Öckinger et al. 2012). Yet, these patterns have not been evaluated in a restoration context, highlighting the need to understand how the spatial configuration of restored habitat influences the diversity of species.

In this study, we answer for the first time the SLOSS question in a restoration context: does restoring a single large habitat patch or multiple small patches, for a fixed amount of restored area, yield higher plant diversity? We measured taxonomic, phylogenetic, and functional diversity of all naturally recruiting woody species in 52 tree islands (hereafter restored patches) in Jambi Province, Sumatra, Indonesia. We evaluated the SLOSS question separately for all species, for native species, and for native forest species. We refer to restored patches as discrete clusters of planted or naturally regenerating tree islands established within a monoculture plantation to promote forest recovery and increase landscape heterogeneity through active or passive restoration. Thirteen restored patches of each of four sizes (25, 100, 400, and 1,600 m²) were established in conventional oil palm plantations six years before vegetation sampling. From these 52 patches, we generated 750 random single large vs. several small comparisons, ensuring the same total area within each comparison (100, 400, or 1,600 m²). Our findings indicate that restoring several small patches results in greater taxonomic, phylogenetic, and functional diversity of recruiting woody species in comparison to a single large patch of the same total area.

## Materials and methods

### Study experiment

We used data collected in the EFForTS-BEE experiment (Ecological and Socioeconomic Functions of Tropical Lowland Rainforest Transformation Systems: Biodiversity Enrichment Experiment), which was established in December 2013. Details about the study region, experiment, and sampling methods are described in full in Teuscher et al. (2016), Zemp et al. (2023), and Paterno et al. (2024).

The study area consisted of 1.4 km^2^ of oil palm plantation in the Jambi province, Sumatra, Indonesia. It also included an orchard, some dirt roads, and small buildings. The wider landscape included other land use types, such as fallow (6%), secondary forest (4%), and rubber plantations (2%), present in patches distributed across the landscape (Korol et al., 2021).

The EFForTS-BEE experiment established 52 restored patches, 13 of each of four patch sizes (25 m², 100 m², 400 m², and 1600 m²) with varying levels of planted tree species diversity (0, 1, 2, 3, and 6 species, where zero represents natural regeneration alone). Tree diversity and composition of the restored patches were initally assigned according to a random partition design (Bell et al. 2009), with up to six native tree species planted. The number of trees planted varied depending on the plot size, with six trees on the 25 m² plots, 25 trees on the 100 m² plots, 100 trees on the 400 m² plots, and 400 trees on the 1,600 m² plots. Restored patches were established within the oil palm plantation, maintaining a minimum distance of 85 m between them, and in a block randomly assigned to a treatment (see Paterno et al., 2024; Fig. 1). In 2020, a comprehensive survey was conducted to record all naturally regenerating woody plants—including trees, shrubs, lianas, and bamboos—taller than 130 cm, totaling 58 plant species, with trees planted as part of experimental treatments excluded from the dataset. This dataset provides a complete census of all recruiting woody species in each of the restored patches. Details about the taxonomic classification, phylogenetic relationships, and functional traits of all plant species can be found in Paterno et al. (2024).

### Creation of single large vs. several small combinations

We assessed whether, for a given total restored area (i.e., 100, 400, or 1,600 m²), a single large restored patch recruited greater taxonomic, phylogenetic, and functional diversity of woody species than several smaller restored patches of varying sizes. We did this by generating random comparisons of single large vs. several small patches, ensuring that the total area of the single large patch and the several smaller ones was the same within each combination. We generated 250 comparisons of single large vs. several small patches for each of three total areas—100, 400, and 1,600 m²—for a total of 750 comparisons. We included in our analysis all 52 experimental restored patches, including those without any recruiting woody species (3 patches of 25 m^2^ when considering all, and forest species; 9 patches of 25 m^2^, 4 patches of 100 m^2^, and 4 patches of 400 m^2^ when considering only native forest species). Sets of several small patches could include any patch that was smaller than the single large ones, with the smaller patches summing to the same total area as the large one (Fig. 2; Supplementary material S1). We could not limit the analyses to combinations with only equal-sized several small patches because there were only 13 patches of each size, i.e., there were not enough 25 m² patches to form sets totaling 400 m² or 1,600 m², and there were not enough 100 m² patches to form sets of several small patches totalling 1,600 m². Thus, for each comparison for the 100-m² total area, we randomly selected one 100-m² single large patch and a set of smaller patches comprised of four 25-m² patches. For the 400-m² total area, we randomly selected one 400-m² single large patch and a set of smaller patches that could be comprised of four 100-m² patches or a combination of 25-m² and 100-m² patches. For the 1,600-m², we randomly selected one 1,600-m² single large patch and a set of smaller patches that could be comprised of four 400-m² patches or any combination of patch sizes summing to 1,600 m².

**Figure 2.**
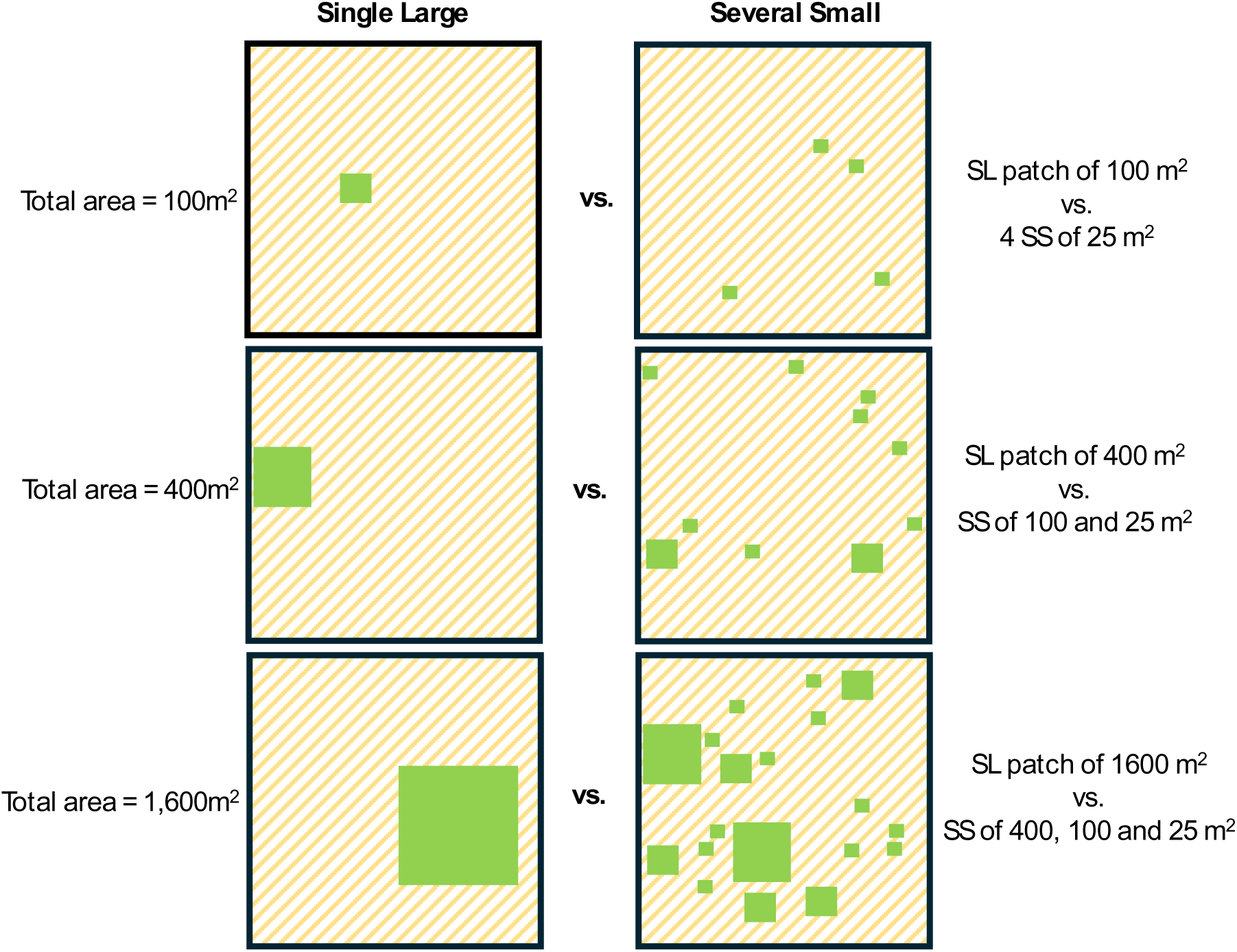
Examples of random selections of Single Large (SL) and Several Small (SS) combinations, where the set of several smaller patches sums to the same total area as the single large patch in each combination.

In all cases, within each combination of several small patches, no individual patch was included more than once, because we were evaluating total diversity (described below) for the species accumulated across a given habitat area equivalent to the single large patch. Repeating individual smaller patches in a given combination would bias results in favour of the single large patch, as the mechanisms by which several smaller patches could support higher species diversity (e.g., greater environmental heterogeneity) would not be fully represented. Finally, although there were only 13 patches of each size, we performed post-hoc analyses—after we had uncovered a general Several Small > Single Large pattern—to compare the set of 13 25-m² patches (totalling 325 m²) to the 400-m² and 1,600-m² single large patches (see Supplementary Materials S4-S6). Although this does not control for habitat amount and is therefore biased in favor of larger patches, the results may help to understand the value of very small (25-m² patches) patches in a restoration context.

### Taxonomic diversity

Taxonomic diversity has been widely used in conservation research (Simberloff, 1970; Tilman et al., 2014). For each single-large-vs.-several-small combination, we measured taxonomic diversity as the effective number of species in the single large patch and in the set of several smaller patches, combining species and their abundances across that set of smaller patches. Effective numbers of species were calculated in three ways: (1) as species richness, (2) as the exponential of the Shannon index, and (3) as the inverse of the Simpson concentration index. These measures correspond to Hill diversities of q-orders 0, 1, and 2, respectively (Chao et al. 2021), which progressively increase the weight given to more abundant species. Specifically, q = 0 reflects species richness and weighs all species equally; q = 1 weighs species proportionally to their relative abundances; and q = 2 gives more weight to abundant species, diminishing the contribution of locally rare species. This approach allows for consideration of both numbers of species and evenness for q = 2 and q = 3, but with a common unit of measurement across all diversity measures, i.e. the effective number of equally abundant species (taxonomic diversity). We used the R package iNEXT.3D 1.0.11 for these calculations (Chao et al., 2021).

### Phylogeny and phylogenetic diversity

Phylogenetic diversity applies greater weight to more unique species, i.e., those that are more phylogenetically distinct (Redding & Mooers, 2006; Winter et al., 2013). We calculated phylogenetic diversity as described above for taxonomic diversity, except here we used the species identities from the single large patch or accumulated across the set of several smaller patches, and the phylogenetic distances among them. Thus, each of our measures of phylogenetic diversity can be interpreted as the effective number of equally-divergent lineages across all several small patches or in the single large patch. For these calculations, we used the pairwise phylogenetic distance matrix that Paterno et al. (2024) constructed with all 58 recruiting species. The authors used the mega-tree provided by Jin & Qian (2019), containing the global angiosperm phylogenetic tree developed by Smith & Brown (2018) and the pteridophyte tree from data in Chave et al. (2009).

### Plant functional traits and functional diversity

Functional diversity (or diversity of species traits in a community) captures the variation in species’ ecological functions (McGill et al., 2006). We calculated functional diversity as described above, for taxonomic diversity, except here we used the species identities and their trait dissimilarities. Thus, each of our measures of functional diversity can be interpreted as the effective number of equally-distinct functional species across all several smaller patches or in the single large patch for each combination. Recruiting species were classified by plant origin as alien, native, or endemic; plant dispersal type as anemochory or zoochory; plant habitat type as forest or open; and plant life form as tree, liana, treelet or shrub. Continuous traits included maximum height (1.7–60 m), specific leaf area (6.4–50.3 mg), and wood density (0.1–0.8 cm^3^). Trait data were missing for a small fraction of species. Missing values were imputed using Phylopars, a multivariate phylogenetic approach for trait imputation (Goolsby et al. 2017; see details in Paterno et al. 2024). We calculated trait dissimilarities using a multi-trait dissimilarity matrix applying a modified Gower distance (Bello et al., 2021).

## Analyses

We used linear models to test whether single large restored patches in oil palm plantations have higher plant taxonomic, phylogenetic, and functional diversity than sets of several smaller patches, given the same cumulative total area. Because single large patches or combinations of several small patches could be included more than once in the models, the assumption of independence of model residuals could be violated. We could not include the ID of the large patch or the combination of small patches as random effects due to model convergence problems. However, the residual values clustered by each large patch ID did not show any evident patterns in the residuals.

The predictor variables were the total area of the single large patch and the set of several smaller patches in each combination, treated as a factor with three levels (100 m², 400 m², 1,600 m²) and the categories single large or several small. We included an interaction term between the total area and the SLOSS type (Single Large or Several Small) to test whether the effect of single large or several small is moderated by total habitat area. Because sets of several smaller patches had the same cumulative area as the single large patch in a given combination, and samples represent complete censuses of each patch, we did not need to account for sampling effort bias in our analysis. We repeated the analyses described above three times: once for all sampled species, once for native species only, and once for native forest-associated species only. Thus, in total we ran 27 linear models (nine response variables × three data subsets). Model assumptions were checked through visual inspection of residuals and statistical tests using the R package DHARMa (Hartig & Hartig, 2017).

Our conclusions were similar whether we used Hill numbers of order 0, 1, or 2; therefore, for simplicity we focus on the results for species richness (order 0) in the main results. See Supplementary Material S2,S3 for the rest of the results. Finally, we calculated the relative importance of each predictor in explaining variation in taxonomic, functional, and phylogenetic species richness of all species, native species, and forest native species. This was calculated based on the lmg metric from the relimp package (Firth, 2016), which decomposes the model’s R² by averaging the additional contribution of each predictor across all possible orderings of the predictors.

## Results

For the 39 single large patches (i.e., the 100-m^2^, 400-m^2^, and 1,600-m^2^ single patches), we found an average of 8.06 (range: 2–18) recruiting woody species, 7.05 (range: 2–16) native species, and 2.72 (range: 0–7) native forest species. In contrast, for the 750 sets of several smaller patches, we found an average of 12.08 (range: 1–27) recruiting woody species, 11.37 (range: 1–24) native species, and 7.60 (range: 0–12) native forest species.

Several small restored patches generally contained greater taxonomic, phylogenetic, and functional diversity of recruiting woody species than a single large restored patch of the same total area (Figs. 3, S2, S3, S7–S9). Furthermore, this pattern was consistent across species diversity comparisons for all species (Fig. 3a–c), native species (Fig. 3d–f), and native forest species (Fig. 3g–i). We also found a significant interaction between SLOSS type (Single Large vs. Several Small) and total area, indicating in most cases that the positive effect of several smaller restored patches on species diversity became stronger as total area, i.e., the size of the single large patch or the sum of the smaller patches, increased.

**Figure 3.**
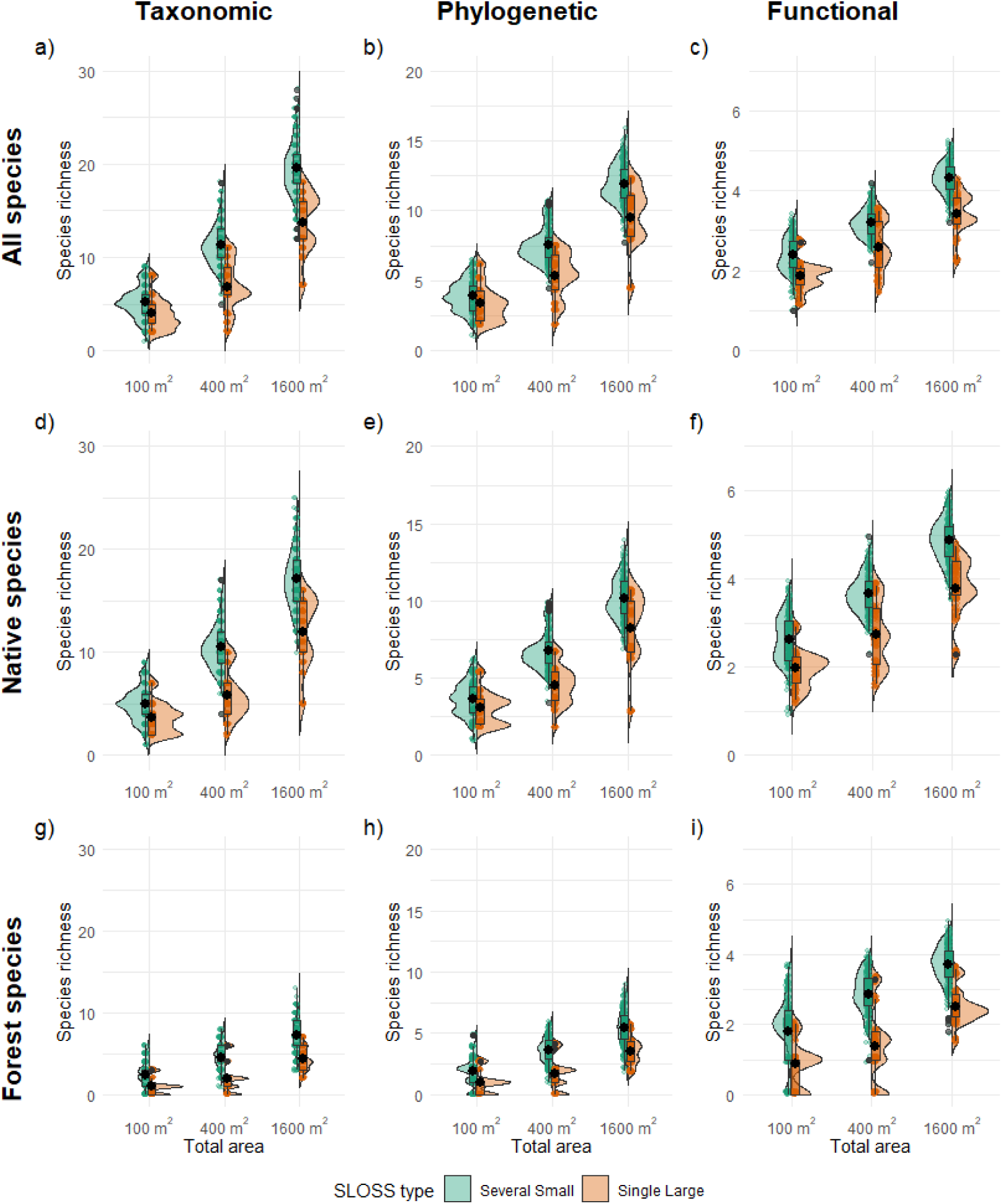
Several small restored patches of varying sizes (green) have significantly higher taxonomic, functional, and phylogenetic diversity of recruiting woody species (excluding planted trees) than a single large restored patch (orange), for the same total restored area. Restored patches are in conventional oil palm plantations. Boxplots show the first and third quartiles, black points and vertical lines indicate the mean and standard error of species diversity for each restoration type and total area combination, and grey points are outliers. Sets of several smaller patches could include patches of any size smaller than the single large patch, summing to its area (“Total area”). Shown are results for species richness; results for the exponential of the Shannon index and the Simpson concentration index are in Figs. S2–S3.

As expected, species diversity increased with total area in all cases, and total area was typically a stronger predictor than SLOSS type and the area × SLOSS interaction (Figs. S10–S12). Total area was the most important variable in 19 of the 27 analyses, whereas SLOSS type was the most important variable in eight of the 27 analyses. These eight cases included: functional diversity of all species, and taxonomic and functional diversity of native species when using the exponential of the Shannon index; taxonomic and functional diversity of all species; and taxonomic, phylogenetic, and functional diversity of native species when using the inverse of the Simpson concentration index.

## Discussion

This is the first study to assess whether a single large patch or several smaller patches, for a fixed total restored area, contains higher taxonomic, phylogenetic, and functional recruiting plant diversity in oil palm plantations. Our analyses show that several smaller restored patches generally support higher diversity of recruiting woody species than a single large restored patch. This result is consistent across three total areas (100, 400, and 1600 m^2^), three diversity measures (species richness, the exponential the of Shannon entropy, and the inverse of Simpson concentration), and when considering all species, only native species, or only native forest species.

A combination of smaller restored patches—including very small ones (25 m^2^)—promotes higher woody species taxonomic, phylogenetic, and functional diversity than a single large patch. This is consistent with previous studies showing that increasing the number of patches in a restoration context may benefit biodiversity equally or even more than restoring the same amount of habitat by increasing patch size or connecting remnants of habitat (Schultz & Crone 2005; Nicol & Possingham 2010; Anonymous, in press). For instance, Anonymous (in press) found that landscapes with a history of restored forest cover had more diverse bird communities when that forest was subdivided into more forest patches. Our results are also consistent with studies assessing restoration strategies that aim to maximize the growth of restored areas by planting more, but smaller tree patches, for a given area to restore (Eppinga et al., 2023; Michaels et al., 2024). Such studies emphasize that planting several smaller tree patches best balances the probability of survival under disturbances and accelerates species colonization compared to planting a single large patch (Fivash et al., 2022). Planting several small tree patches can also accelerate restoration success by enhancing species diversity at the landscape scale through dispersal and/or autocatalytic nucleation (Eppinga et al., 2023). Therefore, our results underscore the importance of fostering heterogeneous landscapes by restoring several small patches to prevent biotic homogenization and increase species turnover and beta diversity (Holl et al., 2020; Brancalion et al., 2025; Holl et al. 2022; Montaya-Sánchez et al., 2023).

Different ecological processes might explain our findings. First, several small restored patches are spread over a larger spatial extent than a single large patch. This could imply greater soil and microhabitat heterogeneity (Duelli, 1997; Fahrig, 2020; Socolar et al., 2016), and lower distances between patches and to scattered trees (Paterno et al. 2024). This could allow a greater diversity of species to establish across several small patches. Second, competition may be lower across several small patches than in a single large one, as dominant species may not recruit into all of the small patches (Hillebrand et al., 2008). This would allow occurrence of more sub-dominant species in several small patches than in a single large one. Third, the number of species planted in several small patches is likely higher in most of the comparisons because it combines the planted trees of all smaller patches in a combination. Higher mean diversity of planted trees in several smaller patches may have led to higher vegetation structural complexity (Kikuchi et al., 2024; Zemp et al., 2019), facilitating the establishment of species with diverse regenerative strategies (Paterno et al. 2016; Paterno et al. 2024). Finally, several small restored forest patches may follow different successional trajectories over time, leading to increased species diversity simply due to ecological drift (Arroyo-Rodríguez et al., 2017; Laurance, 2002; Vellend, 2010). Given the limited number of restored patches, it is not possible to determine which of these processes is/are at play.

That several small restored patches have higher diversity of recruiting woody species than a single large one holds true even when the small patches are very small, i.e., comparing four 25 m² with one 100 m² patch. In fact, in some cases, thirteen 25-m² patches (325 m² in total) even had higher diversity than single patches of 1,600-m^2^ (Figures S4, S5, S6). This suggests that very small restored patches can contribute to the cumulative landscape diversity, in contrast to what is often assumed (Brown & Crone, 2016; Ramirez-Reyes et al., 2016). As 25 m² is the smallest area that could reasonably be described as a restored patch, the results suggests that most of the woody plant species in this system may not have minimum patch size requirements. Overall, the results imply that the benefits of the *number* of restored patches outweigh the costs of reduced patch size.

The diversity benefits of restoring several small patches rather than a single large patch are similar for all woody species groups evaluated, including all species, native species, and native forest species. This implies that most of the native species and even native forest species in the study region may not show minimum patch size effects, at least during the first six years of the experiment. This result challenges the common assumption that specialist species are particularly vulnerable to small patch sizes (Fletcher et al. 2018; Chase et al. 2020), an idea that is largely influenced by studies that confound habitat loss with fragmentation (Fahrig, 2017). However, when controlling for the effect of habitat loss, studies generally find that specialist species are not more negatively affected by fragmentation than generalists (Carrara et al. 2015; Rösch et al. 2015; Fahrig 2020; Riva & Fahrig 2023a; Beard et al. 2025), though there are exceptions (Öckinger et al. 2012).

As expected, species diversity increased strongly with total restored area for both single large and several small restored patches. In other words, the more restored area, the better for biodiversity. This aligns with previous studies showing that species diversity increases with area (Arroyo-Rodríguez et al., 2009; Dendy et al., 2015; Paterno et al., 2024), a consequence of the species-area relationship (Schoener, 1976). As the sum of areas of the largest patches (20,800m^2^) was much greater than the total area of any of the smaller patches sizes, or even all of them summed together, it is also not surprising that 36% of all observed recruiting tree species were observed only in the largest patches. If we could add many more small patches, bringing the cumulative area to 20,800m^2^ this would, according to the species-area relationship, result in additional species being found in those additional small patches. However, we cannot know whether these additional species would include the same species that were observed only in the largest patches. In other words, the 36% of species that were not observed in the smaller patches may have been absent due to patch size effects or due to the smaller total area of small patches in the experiment. This uncertainty highlights the importance of controlling for total area when evaluating patch size effects. Given this uncertainty, along with our finding of more species in several small than a single large patch, and the strong effect of total restored area, we suggest that in oil palm landscapes any and all opportunities for forest restoration should be taken, with the primary aim of maximizing total restored area to promote diversity at the landscape scale.

We recognize that our results may not hold for other taxa. For example, Riva and Fahrig (2023b) found, in a non-restoration context, that the several small patches are better than a single large one pattern is stronger for plants than for other taxa, particularly amphibians and reptiles. The only other analysis of fragmentation in a restoration context that we are aware of is Anonymous (in press), who found that landscapes with a history of restored forest cover had more diverse bird communities when that forest was subdivided into more forest patches. Overall, we need more empirical SLOSS studies in a restoration context, of different taxa in different environments, especially at larger spatial and temporal scales.

In conclusion, our results indicate that restoring a large number of small forest patches—including very small ones—can be an effective strategy for enhancing plant diversity at the landscape scale when planning restoration actions in oil palm plantations. This pattern holds across all biodiversity metrics considered (including taxonomic, functional, and phylogenetic diversity), as well as three Hill numbers (species richness, the exponential of the Shannon index, and the inverse Simpson index), and when focusing on different subsets of species (all species, native species, and native forest species). Our findings challenge studies advocating for restoration strategies that focus exclusively on enlarging single patches or creating large patches by connecting together existing patches in the landscape. Promoting the restoration of multiple small patches of different sizes across a landscape while maximizing the total area restored can be a promising approach to prevent, halt, and reverse biodiversity loss.

## Supplementary material

**Figure S1.**
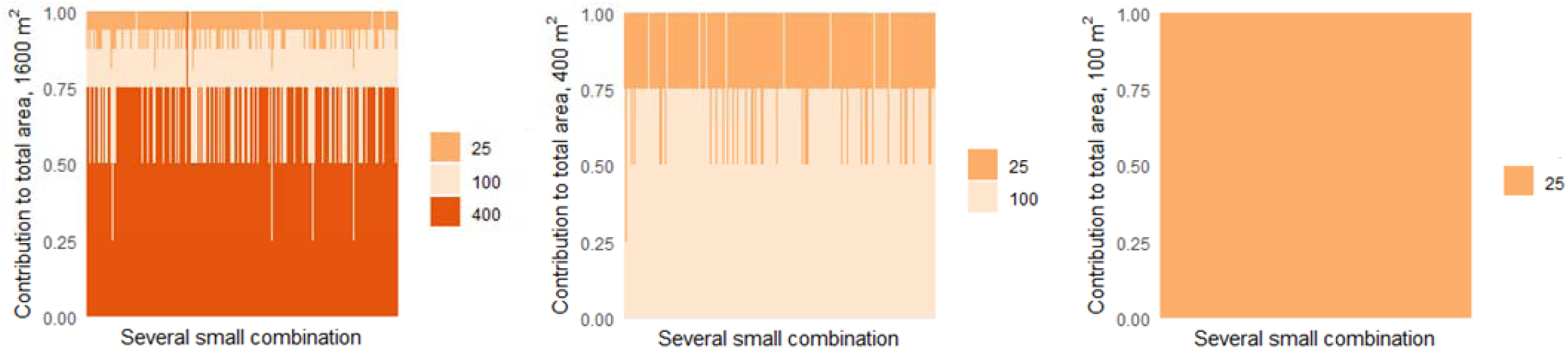
Proportional contribution of patch sizes (25, 100, and 400 m²) to the total area in several-smaller combinations used in the study. Each bar represents a combination of patches summing to 100, 400, or 1600 m². Color intensity corresponds to patch size.

**Figure S2.**
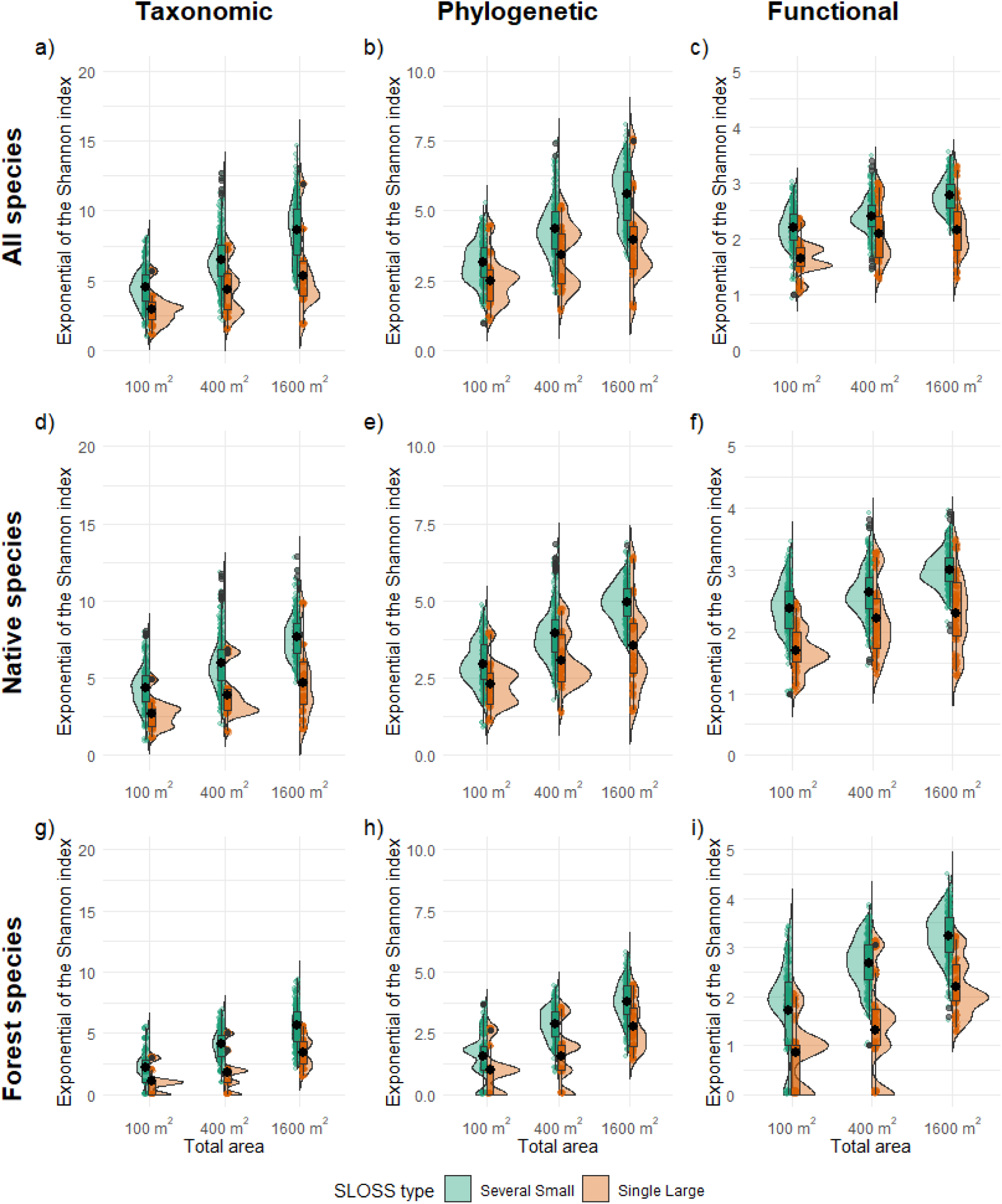
Several small restored patches (green) had significantly higher taxonomic, functional, and phylogenetic diversity of recruiting tree species (excluding planted trees) than a single large restored patch (orange), for the same total restored area. The response is the exponential of the Shannon index. Restored patches were located in conventional oil palm plantations (see Paterno et al. 2024). Boxplots show the first and third quartiles, black points and vertical lines indicate the mean and standard error of species diversity for each restoration type and total area, i.e. area of the single large or the sets of several small patches in a given combination. Grey points represent outliers. Sets of several smaller patches could include patches of any smaller size summing to the total area of the single large patch in a given combination.

**Figure S3.**
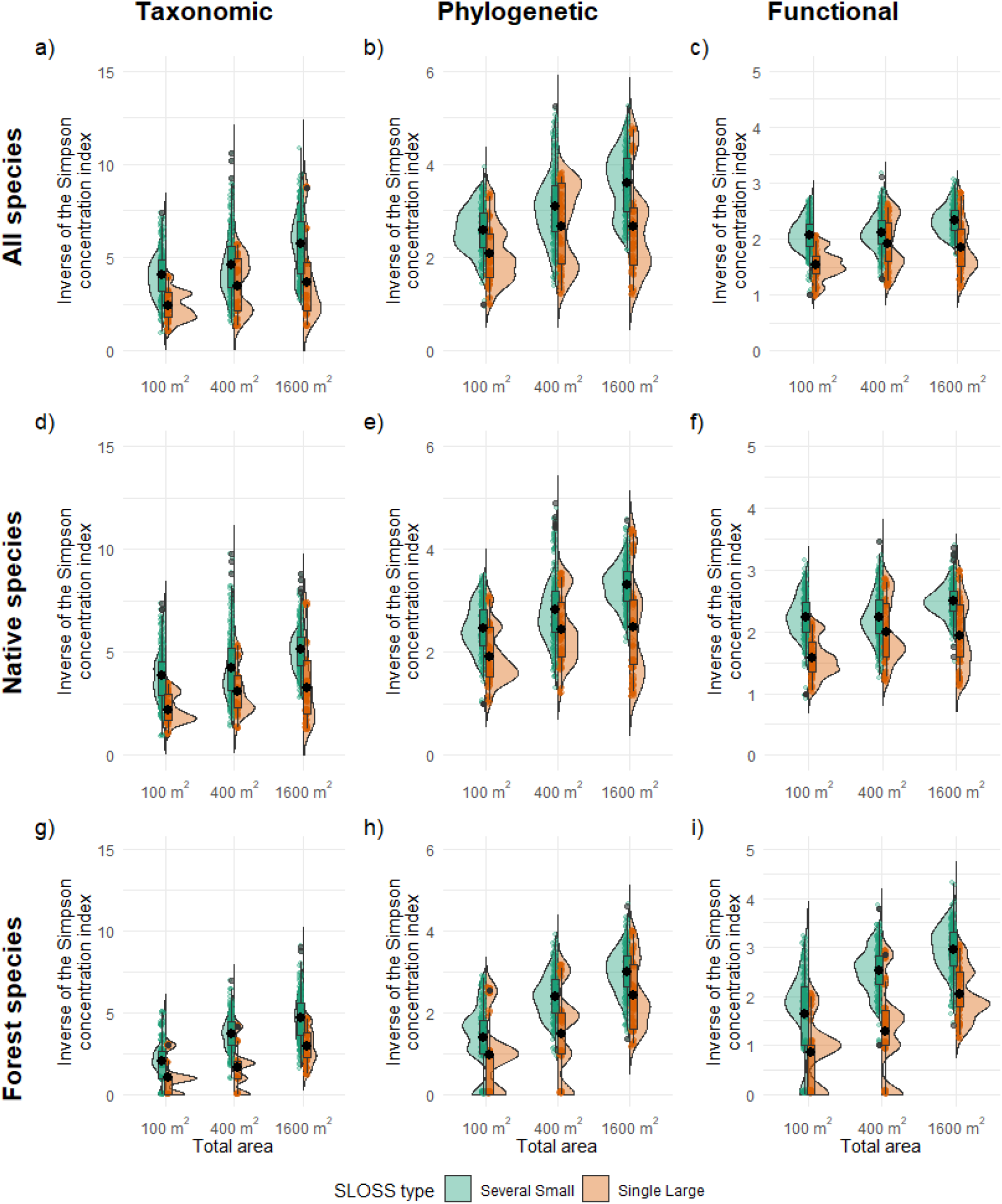
Several small restored patches (green) have significantly higher taxonomic, functional, and phylogenetic diversity of recruiting tree species (excluding planted trees) than a single large restored patch (orange), for the same total restored area when assessing the inverse of the Simpson concentration index. Restored patches are in conventional oil palm plantations. Boxplots show the first and third quartiles, black points and vertical lines indicate the mean and standard error of species diversity for each restoration type and total area combination. and grey points represent outliers. Sets of several smaller patches could include patches of any smaller size summing to the total area.

**Figure S4.**
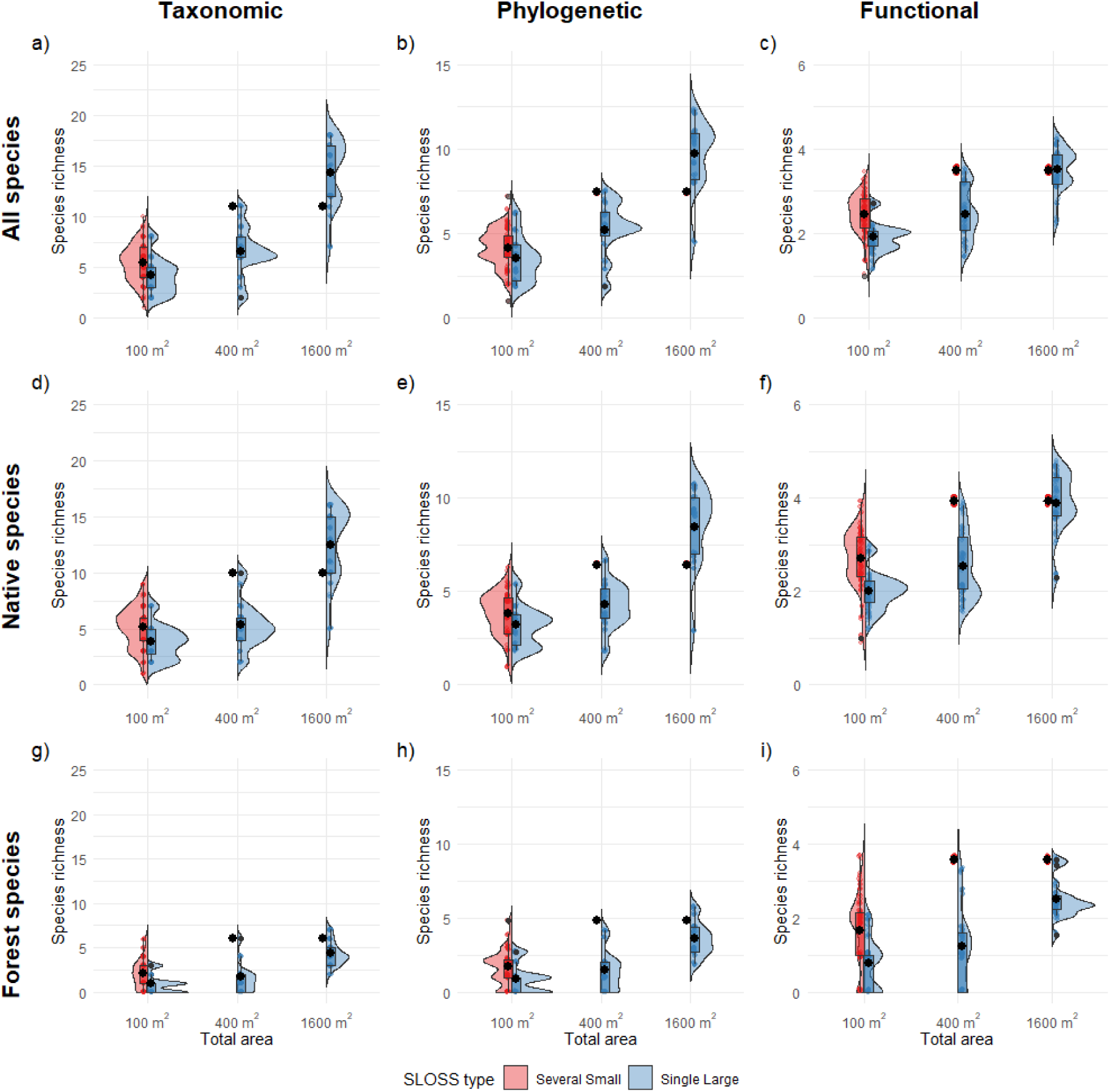
Several small restored patches (red) have significantly higher taxonomic, functional, and phylogenetic diversity of recruiting tree species (excluding planted trees) than a single large restored patch (blue), for the same total restored area when assessing species richness. In this case, several smaller restored patches include only 25-m^2^ patches (max total area 325-m^2^). Restored patches are in conventional oil palm plantations. Boxplots show the first and third quartiles, black points and vertical lines indicate the mean and standard error of species diversity for each restoration type and total area combination. and grey points represent outliers. Sets of several smaller could include patches of any smaller size summing to the total area.

**Figure S5.**
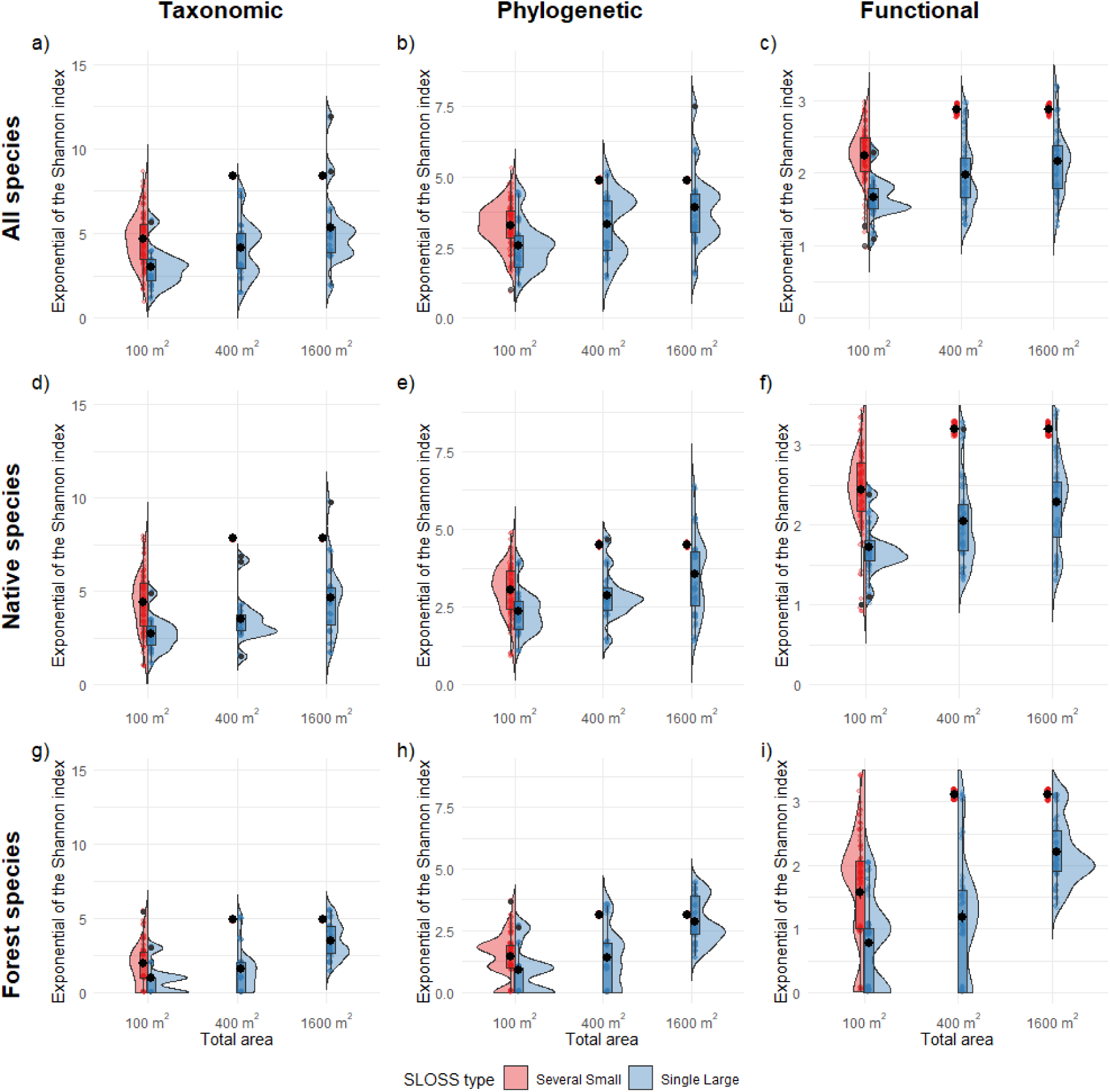
Several small restored patches (red) have significantly higher taxonomic, functional, and phylogenetic diversity of recruiting tree species (excluding planted trees) than a single large restored patch (blue), for the same total restored area when assessing the exponential of the Shannon index. In this case, several smaller restored patches include only 25-m^2^ patches (max total area 325-m^2^). Restored patches are in conventional oil palm plantations. Boxplots show the first and third quartiles, black points and vertical lines indicate the mean and standard error of species diversity for each restoration type and total area combination. and grey points represent outliers. Sets of several smaller patches could include patches of any smaller size summing to the total area.

**Figure S6.**
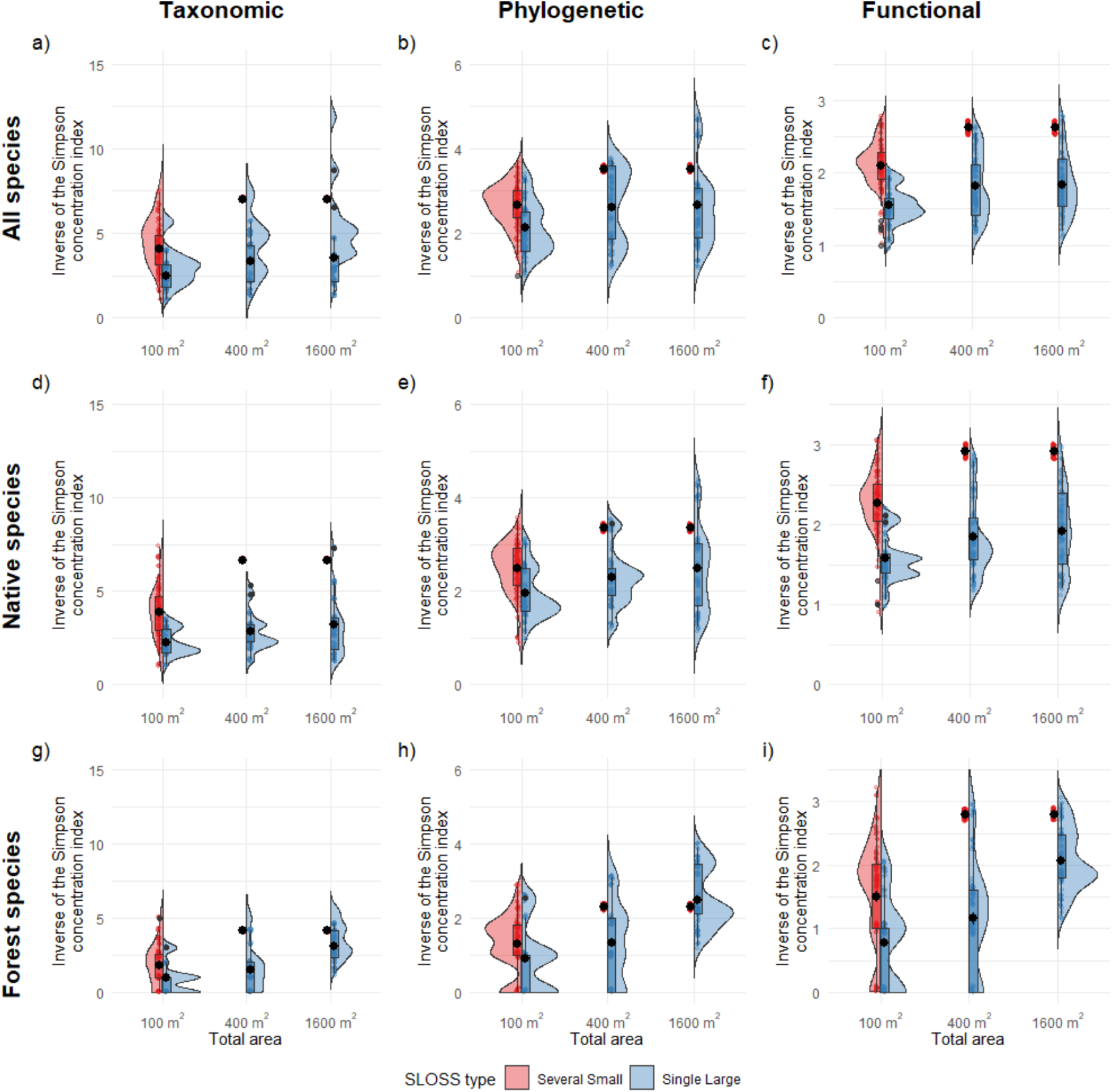
Several small restored patches (red) have significantly higher taxonomic, functional, and phylogenetic diversity of recruiting tree species (excluding planted trees) than a single large restored patch (blue), for the same total restored area when assessing the inverse of the Simpson concentration index. In this case, several smaller restored patches include only 25-m^2^ patches (max total area 325-m^2^). Restored patches are in conventional oil palm plantations. Boxplots show the first and third quartiles, black points and vertical lines indicate the mean and standard error of species diversity for each restoration type and total area combination. and grey points represent outliers. Sets of several smaller could include patches of any smaller size summing to the total area.

**Figure S7.**
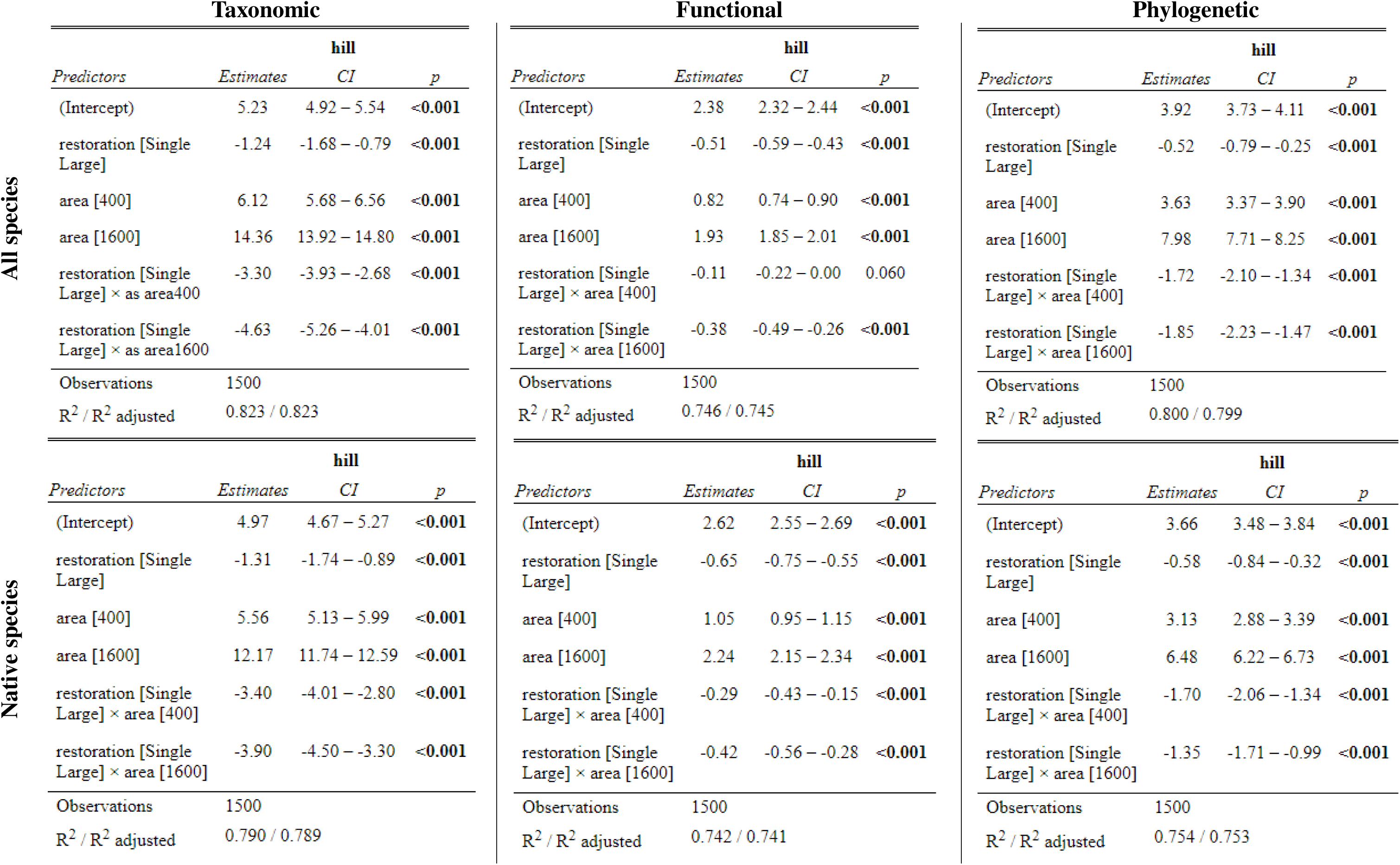
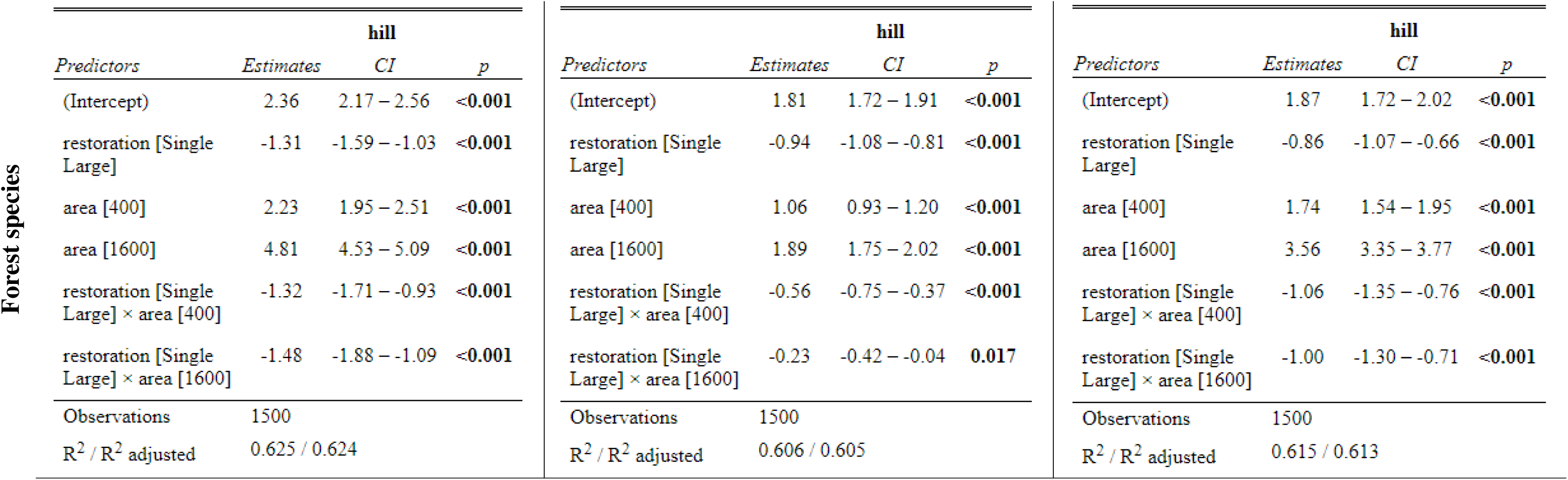
Summary of the linear models assessing the effects of SLOSS type (Single Large vs. Several Small) and total area (100, 400, and 1600 m²) on taxonomic, functional, and phylogenetic species richness (Hill number q = 0) for all species, native species and native forest species. Estimates are based on a Gaussian distribution.

**Figure S8.**
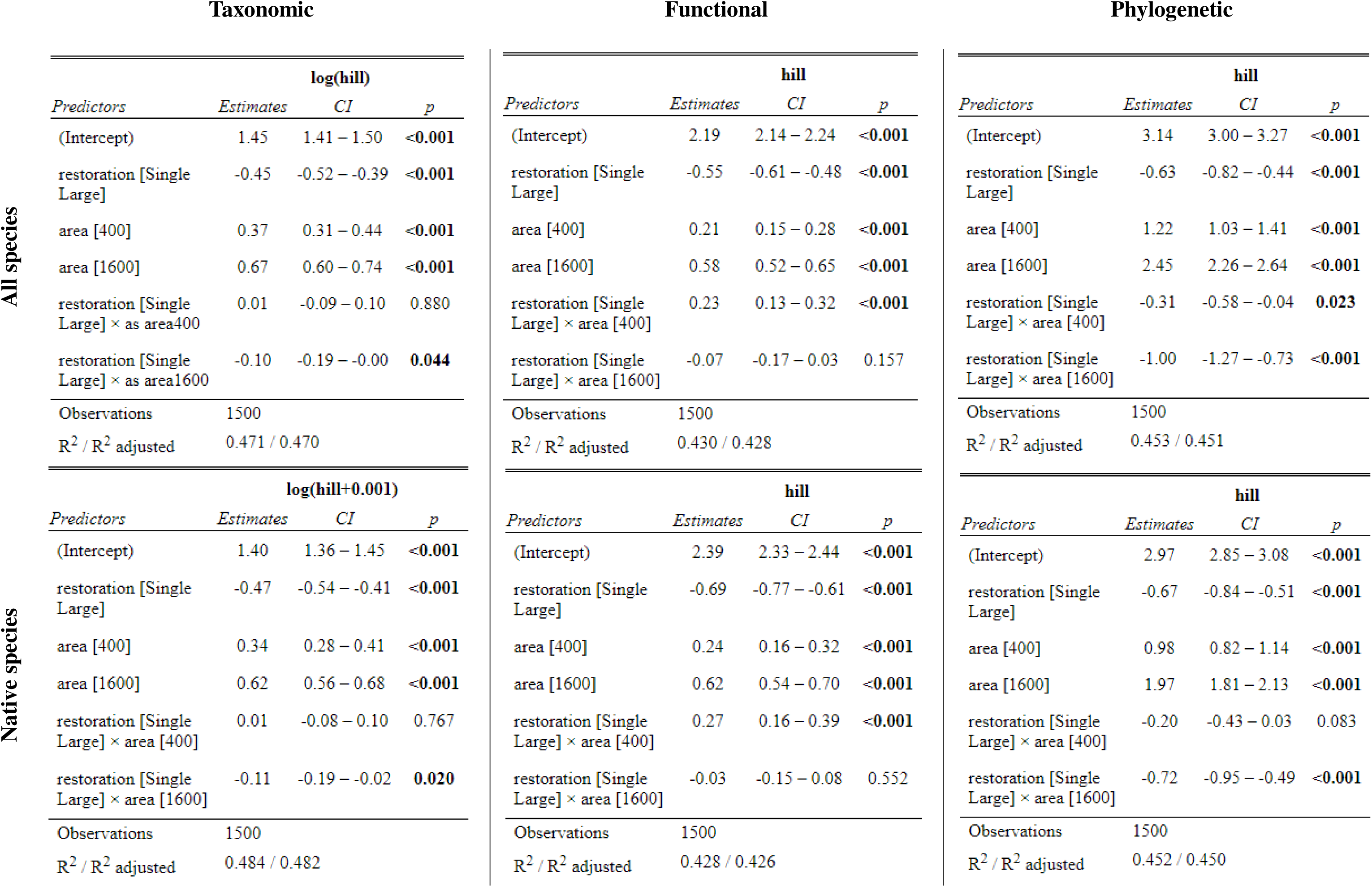
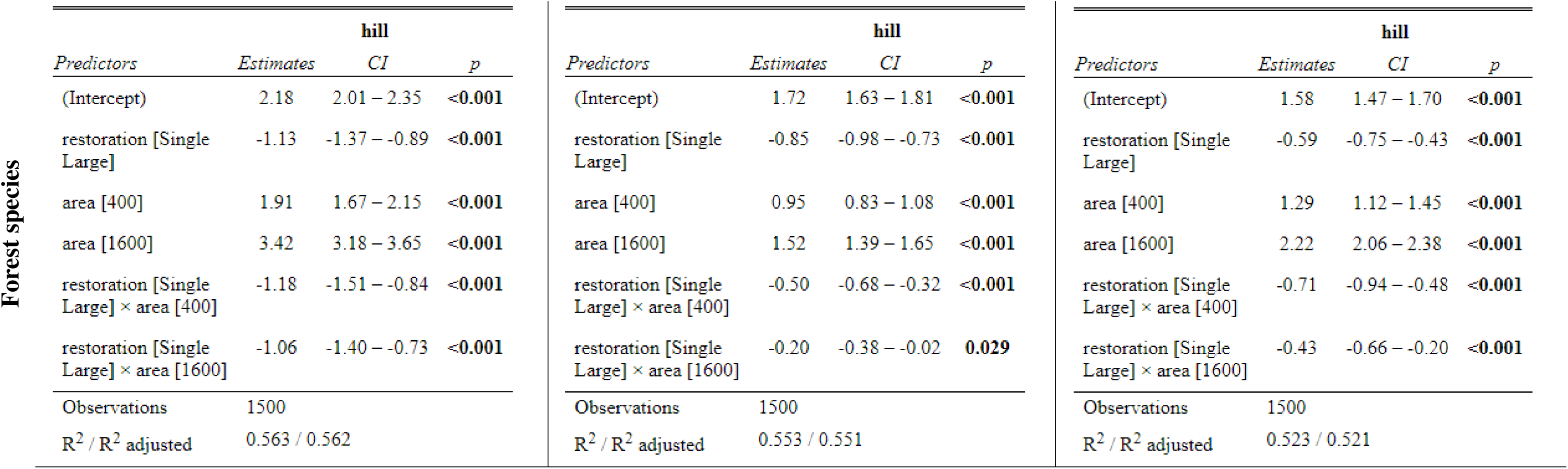
Summary of the linear models assessing the effects of SLOSS type (Single Large vs. Several Small) and total area (100, 400, and 1600 m²) on the taxonomic, functional and phylogenetic exponential of the Shannon index (Hill number q = 1) for all species, native species and native forest species. Estimates are based on a Gaussian distribution.

**Figure S9.**
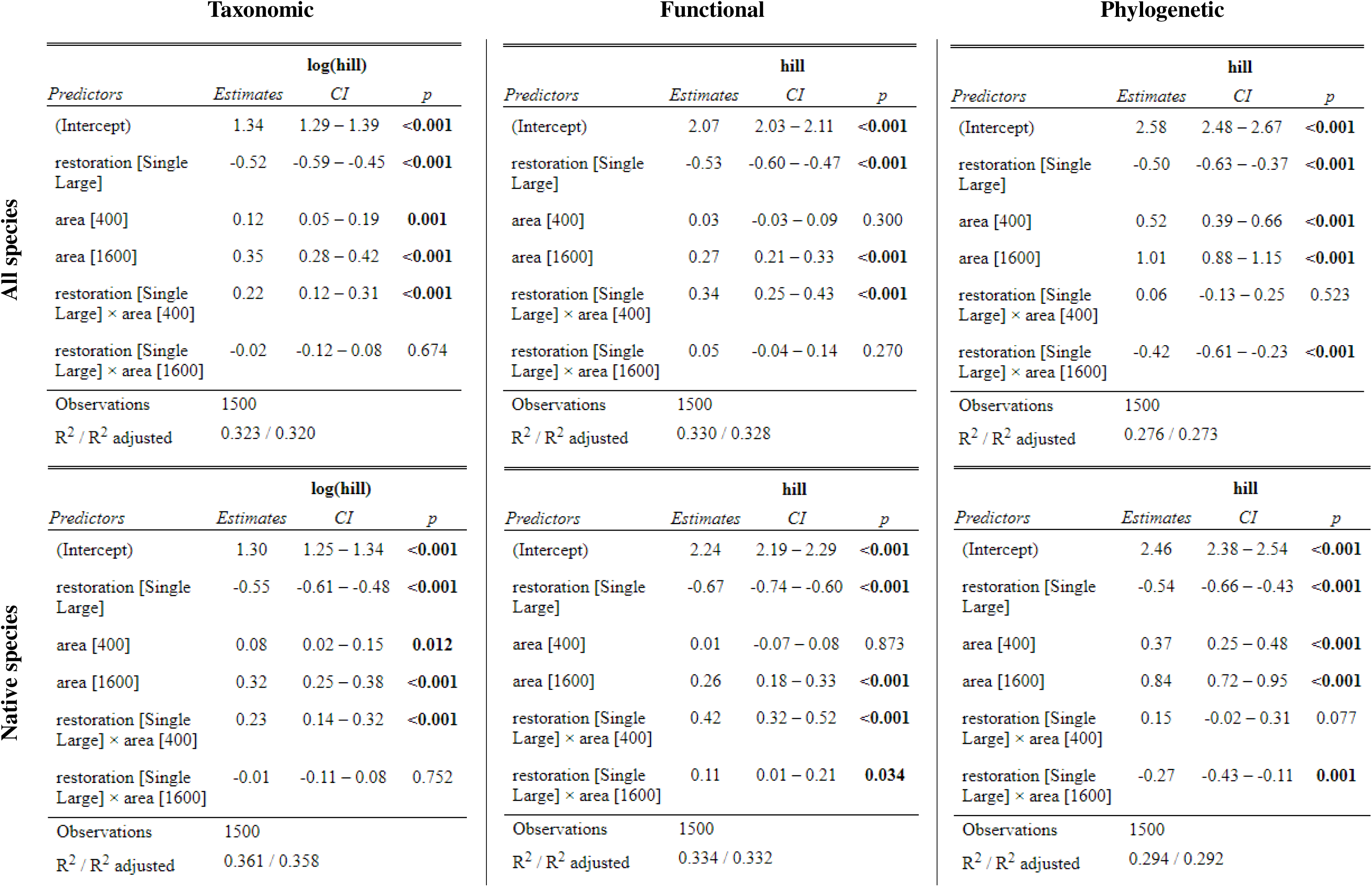
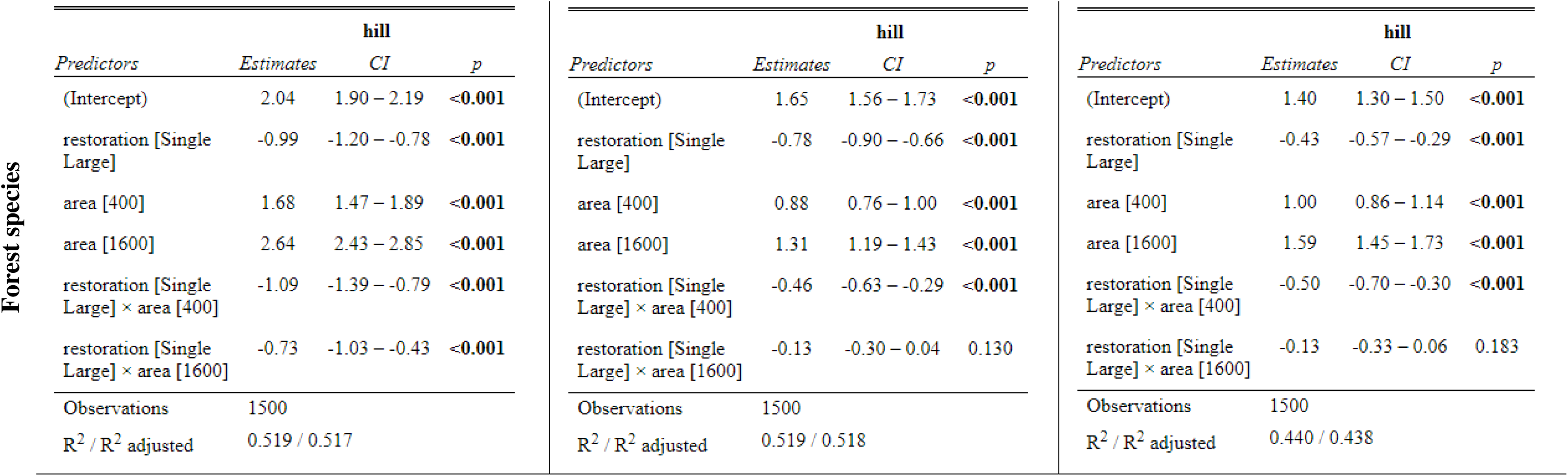
Summary of the linear models assessing the effects of SLOSS type (Single Large vs. Several Small) and total area (100, 400, and 1600 m²) on the taxonomic, functional and phylogenetic Simpson concentration index (Hill number q = 2) for all species, native species and native forest species. Estimates are based on a Gaussian distribution.

**Figure S10.**
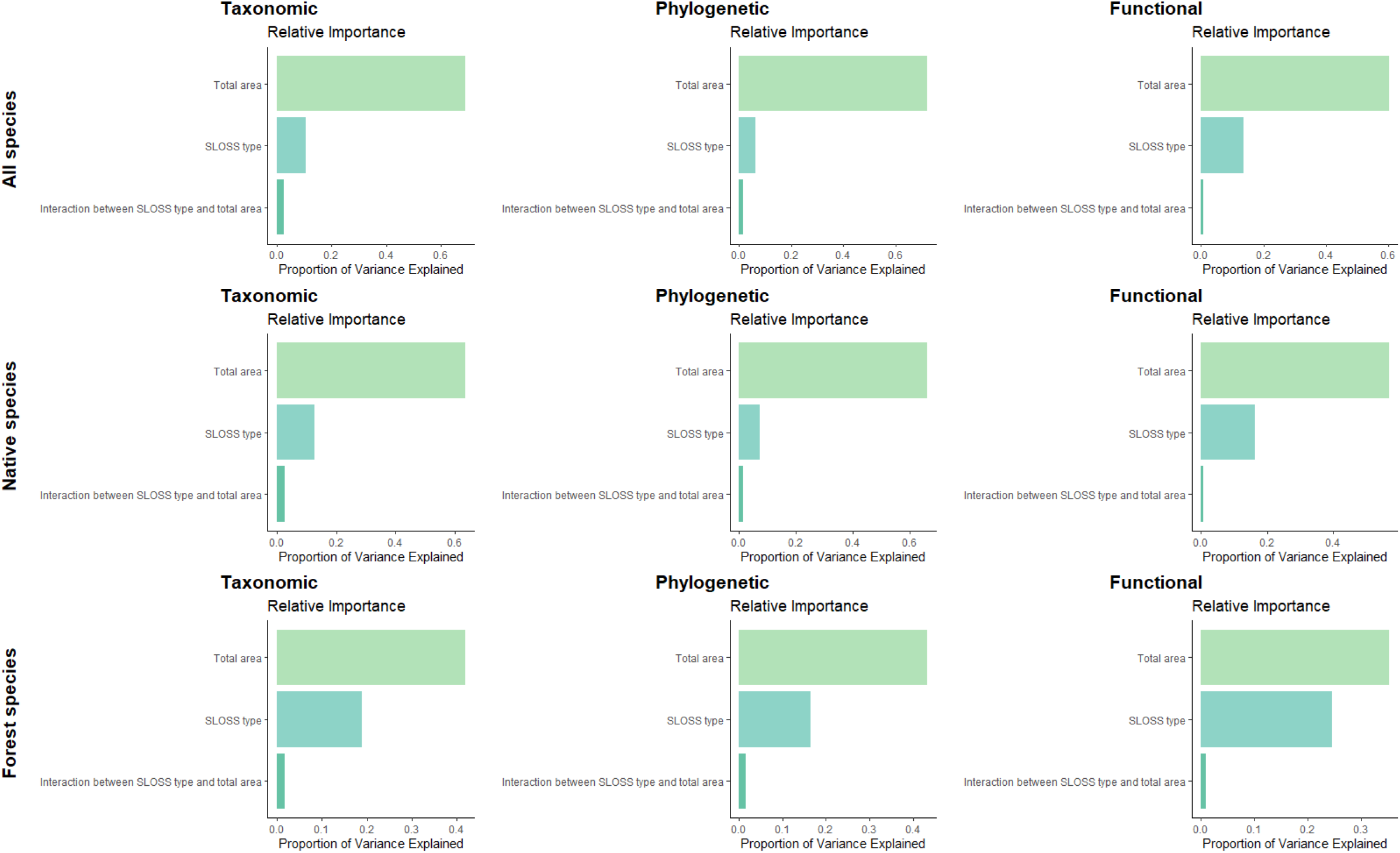
Relative importance of each predictor in explaining variation in taxonomic, functional and phylogenetic species richness (Hill number q = 0) of all species, native species and forest native species. The relative importance is calculated based on the lmg metric from the relimp package in R. The lmg metric decomposes the model’s R² (proportion of variance explained) by averaging the additional contribution of each predictor across all possible orderings. The model includes total area (100, 400, and 1600 m²), SLOSS type (Single Large vs. Several Small), and their interaction. Each bar represents the proportion of variance in species diversity attributed to each predictor or interaction term.

**Figure S11.**
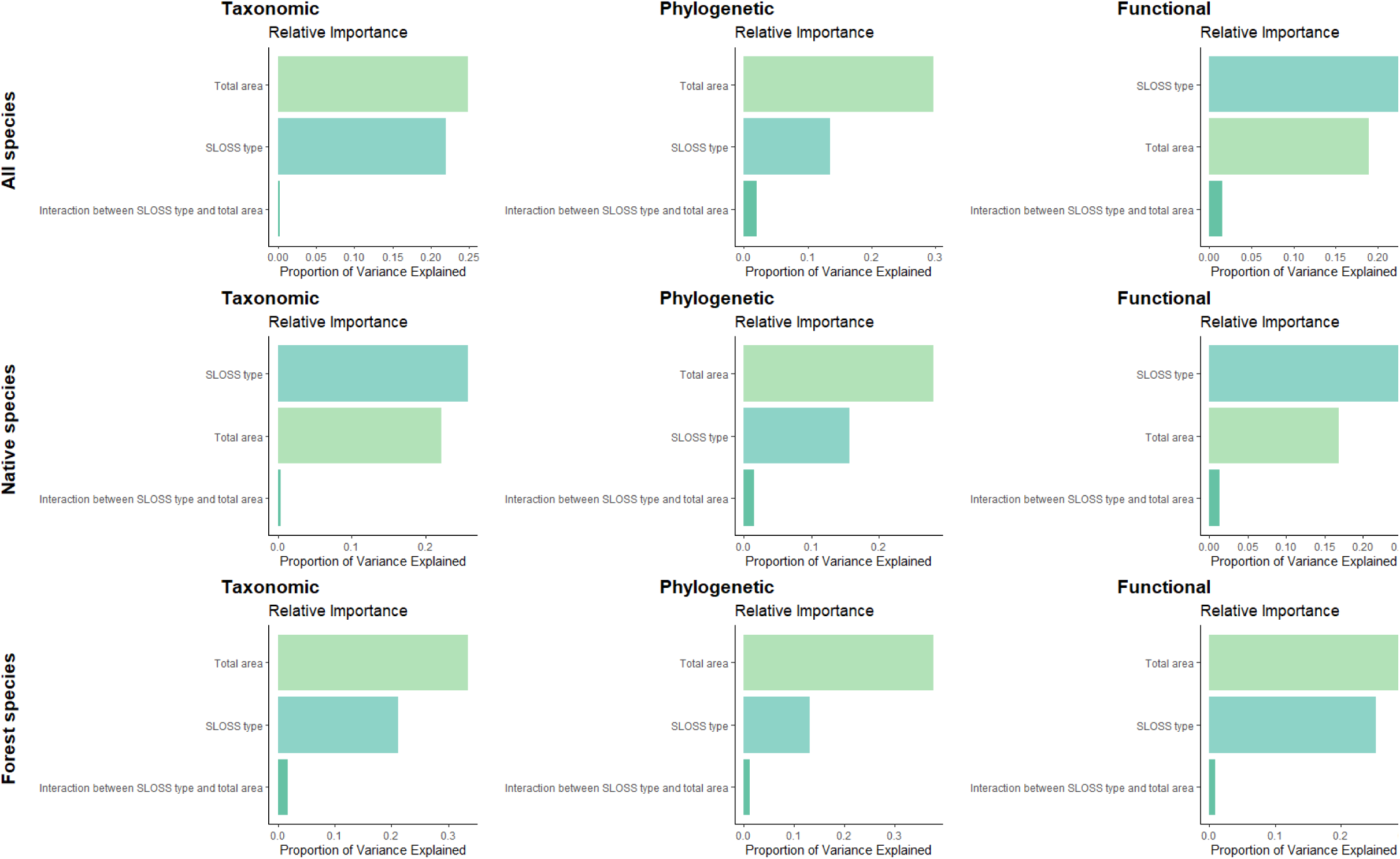
Relative importance of each predictor in explaining variation in taxonomic, functional and phylogenetic exponential of the Shannon index (Hill number q = 1) of all species, native species and forest native species. The relative importance is calculated based on the lmg metric from the relimp package in R. The lmg metric decomposes the model’s R² (proportion of variance explained) by averaging the additional contribution of each predictor across all possible orderings. The model includes total area (100, 400, and 1600 m²), SLOSS type (Single Large vs. Several Small), and their interaction. Each bar represents the proportion of variance in species diversity attributed to each predictor or interaction term.

**Figure S12.**
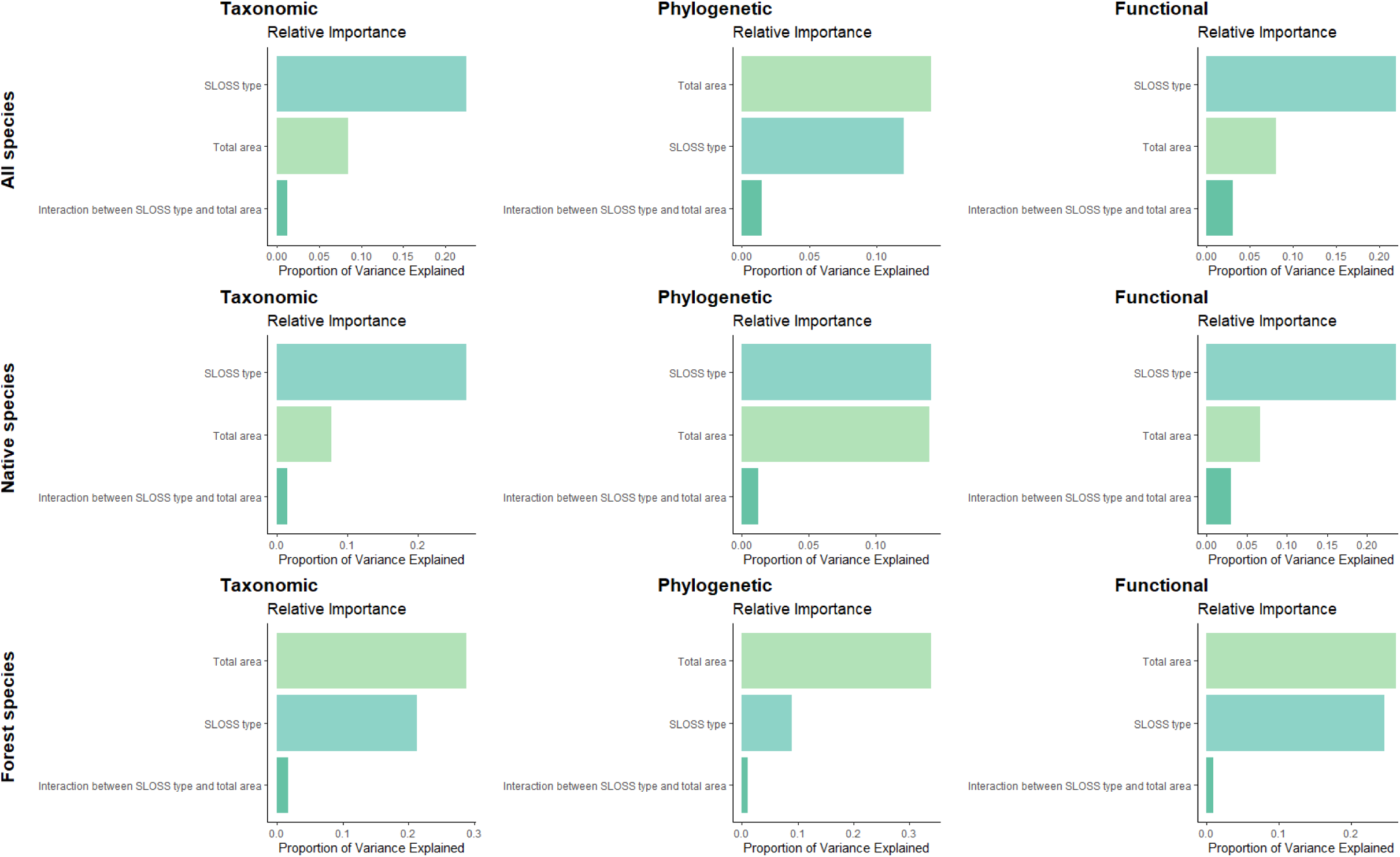
Relative importance of each predictor in explaining variation in taxonomic, functional and phylogenetic Simpson concentration index (Hill number q = 2) of all species, native species and forest native species. The relative importance is calculated based on the lmg metric from the relimp package in R. The lmg metric decomposes the model’s R² (proportion of variance explained) by averaging the additional contribution of each predictor across all possible orderings. The model includes total area (100, 400, and 1600 m²), SLOSS type (Single Large vs. Several Small), and their interaction. Each bar represents the proportion of variance in species diversity attributed to each predictor or interaction term.

## References

1. Antongiovanni, M., Venticinque, E. M., Tambosi, L. R., Matsumoto, M., Metzger, J. P., & Fonseca, C. R. (2022). Restoration priorities for Caatinga dry forests: Landscape resilience, connectivity and biodiversity value. Journal of Applied Ecology, 59(9), 2287–2298.

2. Arroyo-Rodríguez, V., Arasa-Gisbert, R., Arce-Peña, N. P., Cervantes-López, M. J., Cudney-Valenzuela, S. J., Galán-Acedo, C., Hernández-Ruedas, M. A. San-José, M., & Fahrig, L. (2022). The importance of small rainforest patches for biodiversity conservation: a multi-taxonomic assessment. In Biodiversity Islands: Strategies for Conservation in Human-Dominated Environments (pp. 41–60). Springer International Publishing.

3. Arroyo-Rodríguez, V., Melo, F. P. L., Martínez-Ramos, M., Bongers, F., Chazdon, R. L., Meave, J. A., Norden, N., Santos, B. A., Leal, I. R., & M. Tabarelli. (2017). Multiple successional pathways in human-modified tropical landscapes: new insights from forest succession, forest fragmentation and landscape ecology research. Biological Reviews, 92(1), 326–340.

4. Arroyo-Rodríguez, V., Pineda, E., Escobar, F., & Benítez-Malvido, J. (2009). Value of small patches in the conservation of plant-species diversity in highly fragmented rainforest. Conservation Biology, 23(3), 729–739.

5. Banks-Leite, C., Ewers, R. M., Folkard-Tapp, H., & Fraser, A. (2020). Countering the effects of habitat loss, fragmentation, and degradation through habitat restoration. One Earth, 3(6), 672–676.

6. Beard, T., Riva, F., Fahrig, L., & Galán-Acedo, C. (2025). Small patches have high conservation value for primates. Biological Conservation, 311, 111463.

7. Bello, F. de, Botta-Dukát, Z., Lepš, J., & Fibich, P. (2021). Towards a more balanced combination of multiple traits when computing functional differences between species. Methods in Ecology and EvolutionEcol. Evol., 12, 443–448.

8. Brown, L. M., & Crone, E. E. (2016). Minimum area requirements for an at risk butterfly based on movement and demography. Conservation Biology, 30(1), 103–112.

9. Caro, T., Rowe, Z., Berger, J., Wholey, P., & Dobson, A. (2022). An inconvenient misconception: Climate change is not the principal driver of biodiversity loss. Conservation Letters, 15(3), e12868.

10. Carrara, E., Arroyo-Rodríguez, V., Vega-Rivera, J. H., Schondube, J. E., de Freitas, S. M., & Fahrig, L. (2015). Impact of landscape composition and configuration on forest specialist and generalist bird species in the fragmented Lacandona rainforest, Mexico. Biological Conservation, 184, 117–126.

11. Chao, A., Henderson, P. A., Chiu, C., Moyes, F., Hu, K., Dornelas, M., & Magurran, A. E. (2021). Measuring temporal change in alpha diversity: A framework integrating taxonomic, phylogenetic and functional diversity and the iNEXT.3D standardization. Methods in Ecology and Evolution, 12, 1926–1940.

12. Chase, J. M., Blowes, S. A., Knight, T. M., Gerstner, K., & May, F. (2020). Ecosystem decay exacerbates biodiversity loss with habitat loss. Nature, 584(7820), 238–243.

13. Chave, J., Coomes, D., Jansen, S., Lewis, S. L., Swenson, N. G., & Zanne, A. E. (2009). Towards a worldwide wood economics spectrum. Ecology Letters, 12(4), 351–366.

14. Chazdon, R. L., Blüthgen, N., Brancalion, P. H., Heinrich, V., & Bongers, F. (2025). Drivers and benefits of natural regeneration in tropical forests. Nature Reviews Biodiversity, 1–17.

15. Clark, M., Hall, K. R., Martin, D. M., Beaty, B., Lloyd, S., Galgamuwa, G. P., Shirer, R., Zimmerman, C. L., & Shallows, K. M. (2023). Prioritizing restoration sites that improve connectivity in the Appalachian landscape, USA. Conservation Science and Practice, 5(12), e13046.

16. Curtis, P. G., Slay, C. M., Harris, N. L., Tyukavina, A., & Hansen, M. C. (2018). Classifying drivers of global forest loss. Science, 361, 1108–1111.

17. Dendy, J., Cordell, S., Giardina, C. P., Hwang, B., Polloi, E., & Rengulbai, K. (2015). The role of remnant forest patches for habitat restoration in degraded areas of Palau. Restoration Ecology, 23(6), 872–881.

18. Descals, A., Wich, S., Meijaard, E., Gaveau, D. L. A., Peedell, S., & Szantoi, Z. (2021). High-resolution global map of smallholder and industrial closed-canopy oil palm plantations. Earth System Science Data, 13(3), 1211–1231.

19. Diamond, J. M. (1975). The island dilemma: lessons of modern biogeographic studies for the design of natural reserves. Biological Conservation, 7, 129–146.

20. Duelli, P. (1997). Biodiversity evaluation in agricultural landscapes: an approach at two different scales. Agriculture, Ecosystems and Environment, 62, 81–91.

21. Eppinga, M. B., Michaels, T. K., Santos, M. J., & Bever, J. D. (2023). Introducing desirable patches to initiate ecosystem transitions and accelerate ecosystem restoration. Ecological Applications, 33(8), e2910.

22. Fahrig, L. (2017). Ecological responses to habitat fragmentation per se. Annual Review of Ecology, Evolution, and Systematics, 48(1), 1–23.

23. Fahrig, L. (2020). Why do several small patches hold more species than few large patches? Global Ecology and Biogeography, 29, 615–628.

24. Fahrig, L., Watling, J. I., Arnillas, C. A., Arroyo Rodríguez, V., Jörger Hickfang, T., Müller, J., Pereira, H. M., Riva, F., Rösch, V., Seibold, S., Tscharntke, T., & May, F. (2022). Resolving the SLOSS dilemma for biodiversity conservation: a research agenda. Biological Reviews, 97(1), 99–114.

25. Fink, R. D., Lindell, C. A., Morrison, E. B., Zahawi, R. A., & Holl, K. D. (2009). Patch size and tree species influence the number and duration of bird visits in forest restoration plots in southern Costa Rica. Restoration Ecology, 17(4), 479–486.

26. Firth, M. D. (2016). Package ‘relimp’. Relative Contribution of Effects in a Regression Model.

27. Fivash, G. S., van Belzen, J., Temmink, R. J., Didderen, K., Lengkeek, W., van der Heide, T., & Bouma, T. J. (2022). Increasing spatial dispersion in ecosystem restoration mitigates risk in disturbance driven environments. Journal of Applied Ecology, 59(4), 1050–1059.

28. Fletcher Jr, R. J., Didham, R. K., Banks-Leite, C., Barlow, J., Ewers, R. M., Rosindell, J., & Al., E. (2018). Is habitat fragmentation good for biodiversity? Biological Conservation, 226, 9–15.

29. Gann, G. D., McDonald, T., Walder, B., Aronson, J., Nelson, C. R., Jonson, J., Hallett, J. G., Eisenberg, C., Guariguata, M. R., Liu, J., Hua, F., Echeverría, C., Gonzales, E., & Shaw, N. (2019). International principles and standards for the practice of ecological restoration. Restoration Ecology, 27, S1–S46.

30. Gibson, L., Lee, T. M., Koh, L. P., Brook, B. W., Gardner, T. A., Barlow, J., Peres, C. A., Bradshaw, C. J. A., Laurance, W. F., Lovejoy, T. E., & Sodhi, N. S. (2011). Primary forests are irreplaceable for sustaining tropical biodiversity. Nature, 478, 378–383.

31. Goolsby, E. W., Bruggeman, J., & Ané, C. (2017). Rphylopars: fast multivariate phylogenetic comparative methods for missing data and within species variation. Methods in Ecology and Evolution, 8(1), 22–27.

32. Hartig, F., & Hartig, M. F. (2017). *Package ‘DHARMa.’* R Development Core Team.

33. Hillebrand, H., Bennett, D. M., & Cadotte, M. W. (2008). Consequences of dominance: A review of evenness effects on local and regional ecosystem processes. Ecology, 89, 1510–1520.

34. Jin, Y., & Qian, H. (2019). V.PhyloMaker: An R package that can generate very large phylogenies for vascular plants. Ecography, 42, 1353–1359.

35. Kikuchi, T., Seidel, D., Ehbrecht, M., Zemp, D. C., Brambach, F., Irawan, B., Sundawati, L., Hölscher, D., Kreft, H., & Paterno, G. B. (2024). Combining planting trees and natural regeneration promotes long-term structural complexity in oil palm landscapes. Forest Ecology and Management, 569, 122182.

36. Korol, Y., Khokthong, W., Zemp, C. D., Irawan, B., Kreft, H., & Hölscher, D. (2021). Scattered trees in an oil palm landscape: Density, size and distribution. Global Ecology and Conservation, 28, e01688.

37. Laurance, W. F. (2002). Hyperdynamism in fragmented habitats. Journal of Vegetation Science, 13(4), 595–602.

38. Lindenmayer, D. B., Wood, J., McBurney, L., Blair, D., & Banks, S. C. (2015). Single large versus several small: The SLOSS debate in the context of bird responses to a variable retention logging experiment. Forest Ecology and Management, 339, 1–10.

39. Lövei, G. L., Magura, T., Tothmeresz, B., & Kodobocz, V. (2006). The influence of matrix and edges on species richness patterns of ground beetles (Coleoptera : Carabidae) in habitat islands. Global Ecology and Biogeography, 15, 283–289.

40. McGill, B. J., Enquist, B. J., Weiher, E., & Westoby, M. (2006). Rebuilding community ecology from functional traits. Trends in Ecology & Evolution, 21, 178–185.

41. Michaels, T. K., Eppinga, M. B., & Bever, J. D. (2024). When patches grow themselves: From analogy to autocatalytic processes, the relevance of ecological nucleation for restoration practices. Restoration Ecology, 32(1), e14066.

42. Miller, J. E., Damschen, E. I., Harrison, S. P., & Grace, J. B. (2015). Landscape structure affects specialists but not generalists in naturally fragmented grasslands. Ecology, 96(12), 3323–3331.

43. Newbold, T., Hudson, L. N., Arnell, A. P., Contu, S., De Palma, A., Ferrier, S., Hill, S. L. L., Hoskins, A. J., Lysenko, I., Phillips, H. R. P., Burton, V. J., Chng, C. W. T., Emerson, S., Gao, D., Pask-Hale, G., Hutton, J., Jung, M., Sanchez-Ortiz, K., Simmons, B. I., … Purvis, A. (2016). Has land use pushed terrestrial biodiversity beyond the planetary boundary? A global assessment. Science, 353(6296), 288–291.

44. Neyret, M., Richards, D., Prima, M. C., Etherington, T. R., & Lavorel, S. (2025). One cannot have it all: Trading-off ecosystem services and biodiversity bundles in landscape connectivity restoration. Biological Conservation, 302, 110946.

45. Nicol, S. C., & Possingham, H. P. (2010). Should metapopulation restoration strategies increase patch area or number of patches? Ecological Applications, 20(2), 566–581.

46. Öckinger, E., Bergman, K. O., Franzén, M., Kadlec, T., Krauss, J., Kuussaari, M., Pöyry, J., Smith, H. G., Steffan-Dewenter, I., & R. Bommarco. (2012). The landscape matrix modifies the effect of habitat fragmentation in grassland butterflies. Landscape Ecology, 27, 121–131.

47. Ovaskainen, O. (2002). Long-term persistence of species and the SLOSS problem. Journal of Theoretical Biology, 218(4), 419–433.

48. Paterno, G. B., Brambach, F., Guerrero-Ramírez, N., Zemp, D. C., Cantillo, A. F., Camarretta, N., Moura, C., Gailing, C. M., Ballauff, O., Polle, J. A., Schlund, M., Erasmi, S., Iddris, N. A., Khokthong, W., Sundawati, L., Irawan, B., Hölscher, D., & Kreft, H. (2024). Diverse and larger tree islands promote native tree diversity in oil palm landscapes. Science, 386(6723), 795–802.

49. Quinn, J. F., & Harrison, S. P. (1988). Effects of habitat fragmentation and isolation on species richness: evidence from biogeographic patterns. Oecologia, 75(1), 132–140.

50. Ramirez-Reyes, C., Bateman, B. L., & Radeloff, V. C. (2016). Effects of habitat suitability and minimum patch size thresholds on the assessment of landscape connectivity for jaguars in the Sierra Gorda, Mexico. Biological Conservation, 204, 296–305.

51. Redding, D. W., & Mooers, A. Ø. (2006). Incorporating evolutionary measures into conservation prioritization. Conservation Biology, 20(6), 1670–1678.

52. Riva, F., & Fahrig, L. (2022). The disproportionately high value of small patches for biodiversity conservation. Conservation Letters, 15(3), e12881.

53. Riva, F., & Fahrig, L. (2023a). Landscape scale habitat fragmentation is positively related to biodiversity, despite patch scale ecosystem decay. Ecology Letters, 26(2), 268–277.

54. Riva, F., & Fahrig, L. (2023b). Obstruction of biodiversity conservation by minimum patch size criteria. Conservation Biology, 37(5), e14092.

55. Riva, F., Galán Acedo, C., Martin, A. E., & Fahrig, L. (2025). Why we should not assume that habitat fragmentation is generally bad for restoration: a reply to Watts and Hughes (2024). Restoration Ecology, e14385.

56. Rösch, V., Tscharntke, T., Scherber, C., & Batáry, P. (2015). Biodiversity conservation across taxa and landscapes requires many small as well as single large habitat fragments. Oecologia, 179(1), 209–222.

57. Schoener, T. W. (1976). The species–area relationship within archipelagoes: models and evidence from island birds. Proceedings of the XVI International Ornithological Congress, 6, 629–642.

58. Schultz, C. B., & Crone, E. E. (2005). Patch size and connectivity thresholds for butterfly habitat restoration. Conservation Biology, 19(3), 887–896.

59. Simberloff, D. S. (1970). Taxonomic diversity of island biotas. Evolution, 24, 23–47.

60. Smith, S. A., & Brown, J. W. (2018). Constructing a broadly inclusive seed plant phylogeny. American Journal of Botany, 105, 302–314.

61. Socolar, J. B., Gilroy, J. J., Kunin, W. E., & Edwards, D. P. (2016). How Should Beta-Diversity Inform Biodiversity Conservation? Trends in Ecology and Evolution, 31(1), 67–80.

62. Stephens, T. (2023). The Kunming–Montreal Global Biodiversity Framework. International Legal Materials, 62(5), 868–887.

63. Struebig, M. J., Lee, J. S., Deere, N. J., Gevaña, D. T., Ingram, D. J., Lwin, N., Nguyen, T., Santika, T., Seaman, D. J. I., Supriatna, J., & Davies, Z. G. (2025). Drivers and solutions to Southeast Asia’s biodiversity crisis. Nature Reviews Biodiversity, 1–18.

64. Teuscher, M., Gérard, A., Brose, U., Buchori, D., Clough, Y., Ehbrecht, M., Hölscher, D., Irawan, B., Sundawati, L., Wollni, M., & Kreft, H. (2016). Experimental biodiversity enrichment in oil-palm-dominated landscapes in Indonesia. Frontiers in Plant Science, 7, 1–15.

65. Tilman, D., Forest, I., & Cowles, J. M. (2014). Biodiversity and ecosystem functioning. Annual Review of Ecology, Evolution, and Systematics, 45(1), 471–493.

66. Vellend, M. (2010). Conceptual synthesis in community ecology. The Quarterly Review of Biology, 85(2), 183–206.

67. Watts, K., & Hughes, S. (2024). Fragmentation impacts may be mixed for conservation but generally bad for restoration. Restoration Ecology, 32(8), e14260.

68. Winter, M., Devictor, V., & Schweiger, O. (2013). Phylogenetic diversity and nature conservation: Where are we? Trends in Ecology and Evolution, 28(4), 199–204.

69. Wintle, B. A., Kujala, H., Whitehead, A., Cameron, A., Veloz, S., Kukkala, A., Moilanene, A., Gordong, A., Lentinia, P. E., Cadenhead, N. C. R., & Bekessy, S. A. (2019). Global synthesis of conservation studies reveals the importance of small habitat patches for biodiversity. Proceedings of the National Academy of Sciences, 116(3), 909–914.

70. Zemp, D. C., Ehbrecht, M., Seidel, D., Ammer, C., Craven, D., Erkelenz, J., Irawan, B., Sundawati, L., Hölscher, D., & Kreft, H. (2019). Mixed-species tree plantings enhance structural complexity in oil palm plantations. Agriculture, Ecosystems & Environment, 283, 106564. 10.1016/j.agee.2019.06.003

71. Zemp, D. C., Guerrero-Ramirez, N., Brambach, F., Darras, K., Grass, I., Potapov, A., Röll, A., Arimond, I., Ballauff, J., Behling, H., Berkelmann, D., Biagioni, S., Buchori, D., Craven, D., Daniel, R., Gailing, O., Ellsäßer, F., Fardiansah, R., Hennings, N., … Kreft, H. (2023). Tree islands enhance biodiversity and functioning in oil palm landscapes. Nature, 618(7964), 316–321.

